# RNA-seq reveals transcriptomic differences in circadian-related genes of the choroid plexus in a preclinical chronic migraine model

**DOI:** 10.1101/2025.06.03.657705

**Authors:** Yohannes W. Woldeamanuel, Chenchen Xia, Sam Ding, Alfred Fonteh, Xianghong Arakaki

## Abstract

**Background:** Migraine patients show choroid plexus (CP) changes, impairing the blood-CSF barrier. The CP regulates circadian rhythms, but links between CP circadian genes and migraine are unexplored.

**Objective:** This study examined CP circadian gene transcriptome changes in a chronic migraine rat model versus controls to identify migraine-related pathways.

**Design:** Chronic migraine model: Sprague Dawley rats (3 females, 3 males) received nitroglycerine (NTG) every other day for 9 days; controls (3 females, 3 males) got saline. CP from the 4th ventricle was collected 2 hours post-final injection for RNAseq.

**Methods:** Migraine Behavior: Von Frey hair tests on days 1, 5, and 9, pre- and 2 hours post-NTG/saline injection, assessed basal and NTG-induced pain thresholds.

RNAseq & Analysis: Differentially expressed genes (p < 0.05, fold change > 1) were identified. GO, KEGG, and Reactome enrichment analyses evaluated circadian gene expression changes.

**Results:**

- NTG group showed reduced basal and NTG-induced pain thresholds on days 1, 5, and 9.
- Females had more upregulated genes (*MT2A*, *SLC7A11*), males upregulated *ZBTB16*, *S100A8*. *SLC7A11*, *SCG2*, *GRIA1* showed inverse regulation (up in females, down in males).
- Circadian gene expression altered: 10 genes upregulated (e.g., *SERPINE1, MAPK9, ATF4*), 13 downregulated (e.g., *PER2, DBP, EZH2*). Sex-specific differences: females (*FBXL12, GPR157*), males (*NKX2-1, ATF4, CLOCK*).
- GO/KEGG analyses revealed significant enrichment of circadian rhythm-related pathways, insulin resistance, and inflammatory response processes, with sex-specific differences: females showed HIF-1 signaling and hemoglobin-related pathways, while males exhibited arachidonic acid and leukotriene production.

**Conclusion:** CP transcriptomics in the rat migraine model revealed sex-specific gene regulation, with females upregulating antioxidant genes (MT2A, SLC7A11) and males upregulating inflammatory factors (ZBTB16, S100A8), alongside circadian disruption (e.g., SERPINE1 upregulated, PER2 downregulated). Pathway analyses indicate enriched circadian rhythms, HIF-1 signaling (females), inflammatory processes (males), lipid metabolism (PPARα), and heme signaling, highlighting sex-specific and circadian targets for migraine therapy.

## Introduction

Circadian rhythm disruptions, often caused by irregular sleep patterns^1^, inconsistent meal times^2^, and inadequate natural light exposure, are linked to migraine^3–7^. The choroid plexus, the primary source of the cerebrospinal fluid (CSF), plays a crucial role due to its significant circadian oscillation amplitude compared to other brain regions^8–11^. Migraine patients exhibit alterations in their choroid plexus^10,12,13^, resulting in compromised integrity of the blood-CSF barrier (BCB)^13^. This, in turn, is linked to changes in CSF composition, including elevated levels of cell adhesion molecules, specifically soluble vascular cell adhesion molecule-1 (sVCAM-1)^14^. A recent study demonstrated that the trigeminal ganglion has direct contact with CSF^15^. Furthermore, CSF proteomic alterations following cortical spreading depression, associated with the pathophysiology of migraine aura, can activate trigeminal pathways^15^.

While the choroid plexus’s role in circadian rhythms is recognized^9,11^, research on its gene expression related to the regulatory mechanisms in migraine pathology is lacking. This knowledge gap suggests a promising avenue for investigation, as exploring these connections may reveal new insights into migraine mechanisms and treatments. Historically, attention was primarily directed towards the blood-brain barrier (BBB) that forms the neurovascular unit, often leaving the BCB of the choroid plexus underestimated in brain conditions^16^. However, this perspective is now changing due to advances in molecular and imaging technologies, and the choroid plexus’s significance is gaining recognition for its unique role in brain conditions^16,17^.

Unlike the BBB, BCB capillaries do not have the tight junctions that make the BBB highly selective and semipermeable^17^. Instead, they possess gap junctions and are fenestrated, enhancing their permeability to larger macromolecules^17^. The openness of these junctions varies throughout the day due to circadian rhythms, affecting substance transport across this barrier. The BCB monitors CSF composition and triggers rapid immune responses during neurological challenges^17,18^. The BCB contains specialized immune cells, including epiplexus or Kolmer cells (macrophages), T cells, and dendritic cells situated on the CSF-facing apical side of the epithelium^17,18^. These cells are essential for immunosurveillance^17,18^. Thus, the choroid plexus maintains the brain environment by clearing metabolites and neurotoxins. Immune cells bypass the restrictive BBB by using the more permissive BCB to access the brain parenchyma^18^. This mechanism illustrates how the choroid plexus acts as a critical hub for immune cell trafficking and neuroimmune interactions, responding effectively to brain conditions^18^. Neuroimmune crosstalk significantly contributes to the development of chronic migraine^19^. During this process, neuropeptides are released, which attract immune cells that, in turn, secrete cytokines^19^. This activation affects trigeminal nociceptors, resulting in both peripheral and central sensitization^19^.

The choroid plexus synthesizes many proteins and growth factors (e.g., glial-derived and brain-derived neurotrophic factors) that significantly influence brain architecture, neuronal function, neurogenesis, and regulatory mechanisms essential for maintaining brain homeostasis^20^. Circadian rhythms in choroid plexus epithelial cells exhibit sexual dimorphism influenced by estrogen^11^. Both sexes express standard circadian clock genes such as Per2 and Cry2, but Per1, Bmal1, and Clock oscillations are exclusive to females^11,21^. A thorough understanding of transcriptome-level alterations in the choroid plexus can reveal gene pathways associated with migraine regulation. This study comprehensively analyzed gene expression in the choroid plexus using a preclinical chronic migraine model to explore the relationships between circadian-related genes and chronic migraine.

## Methods

### Chronic migraine model and sample collection

Twelve 2-3-month-old Sprague Dawley (*Rattus norvegicus*) rats were used in this study. The experimental group, comprising three female and three male rats, received nitroglycerin injections every other day for nine days. In contrast, the control group, also with three females and three males, received saline. All animals were euthanized two hours post-injection, and the CP of the fourth ventricle was collected for RNA sequencing.

### Migraine behavior assessments

To determine mechanical pain thresholds, we performed behavioral evaluations using the von Frey hair test on the periorbital skin on days 1, 5, and 9 of the experiment, both before and two hours after the NTG or saline injection, to assess basal and NTG-induced pain thresholds, respectively.

### RNA sequencing and gene enrichment analysis

The reads were first mapped to the latest UCSC transcript set (rn6)^22^ using Bowtie2 version 2.1.0^23^, and the gene expression level was estimated using RSEM v1.2.15^24^. Differentially expressed genes (DEGs) were identified through the edgeR^25^ program. Genes with p-values less than 0.05 exhibiting 1.5-fold changes were deemed differentially expressed.

DEGs were identified by calculating fold change (FC = Mean(NTG)/Mean(Saline)) from normalized RNAseq data (Transcripts Per Million/TPM) for female-only, male-only, and combined (male and female) analyses. Due to low statistical power (n=3 per group), genes with FC > 1.5 (upregulated) or < 0.67 (downregulated) for females and males, and FC > 1/<1 for the combined analyses, were selected for pathway enrichment in ShinyGO v0.82^26^ (Rattus norvegicus, FDR < 0.2) to perform the GO (Gene Ontology) enrichment analysis, KEGG (Kyoto Encyclopedia of Genes and Genomes)^27^ pathway enrichment analysis, and Reactome.

### Circadian rhythm-related genes

Genes linked to circadian rhythms were detected by searching for “circadian” within protein and pathway descriptions on genecards.org. This method was similar to that used in a recent transcriptomics study of the choroid plexus that examined day-night variations^28^. For the circadian rhythm-associated genes, a 1-fold change was used as a cutoff to have increased sensitivity of differentially expressed genes.

### Statistical Analysis

Gene expression data underwent auto-scaling, meaning it was mean-centered and divided by the standard deviation of each variable for normalization. No data filtering was performed. Differential gene expression was visualized using a volcano plot, which depicted the relationship between gene expression fold change and statistical significance (-log10 p-value) to identify genes with significant expression changes. The clustering results were visualized as a heatmap, which was generated using Euclidean distance as the distance measure and Ward’s linkage clustering algorithm, which minimizes the sum of squares of any two clusters. No statistical analysis was performed on the behavioral data, as this study aimed to qualitatively demonstrate the established model of nitroglycerin (NTG)-induced migraine-like behavior. Consistent with previous findings, NTG administration was observed to lower mechanical thresholds, indicating increased sensitivity.

## Results

### Mechanical Pain Threshold Comparison

Key Findings

- Basal Pain Thresholds: Mechanical pain thresholds decreased progressively over time in the NTG group, compared to the SAL group (Figure 1A).
- NTG-induced Pain Thresholds: Chronic administration of nitroglycerin induces a long-lasting hypersensitivity with reduced mechanical pain thresholds in the NTG group compared to the SAL group (Figure 1B).
- No statistical analysis was performed on the behavioral data, as this study aimed to qualitatively demonstrate the established model of nitroglycerin (NTG)-induced migraine-like behavior. Consistent with previous findings, NTG administration was observed to lower mechanical thresholds, indicating increased sensitivity.

**Figure 1.**
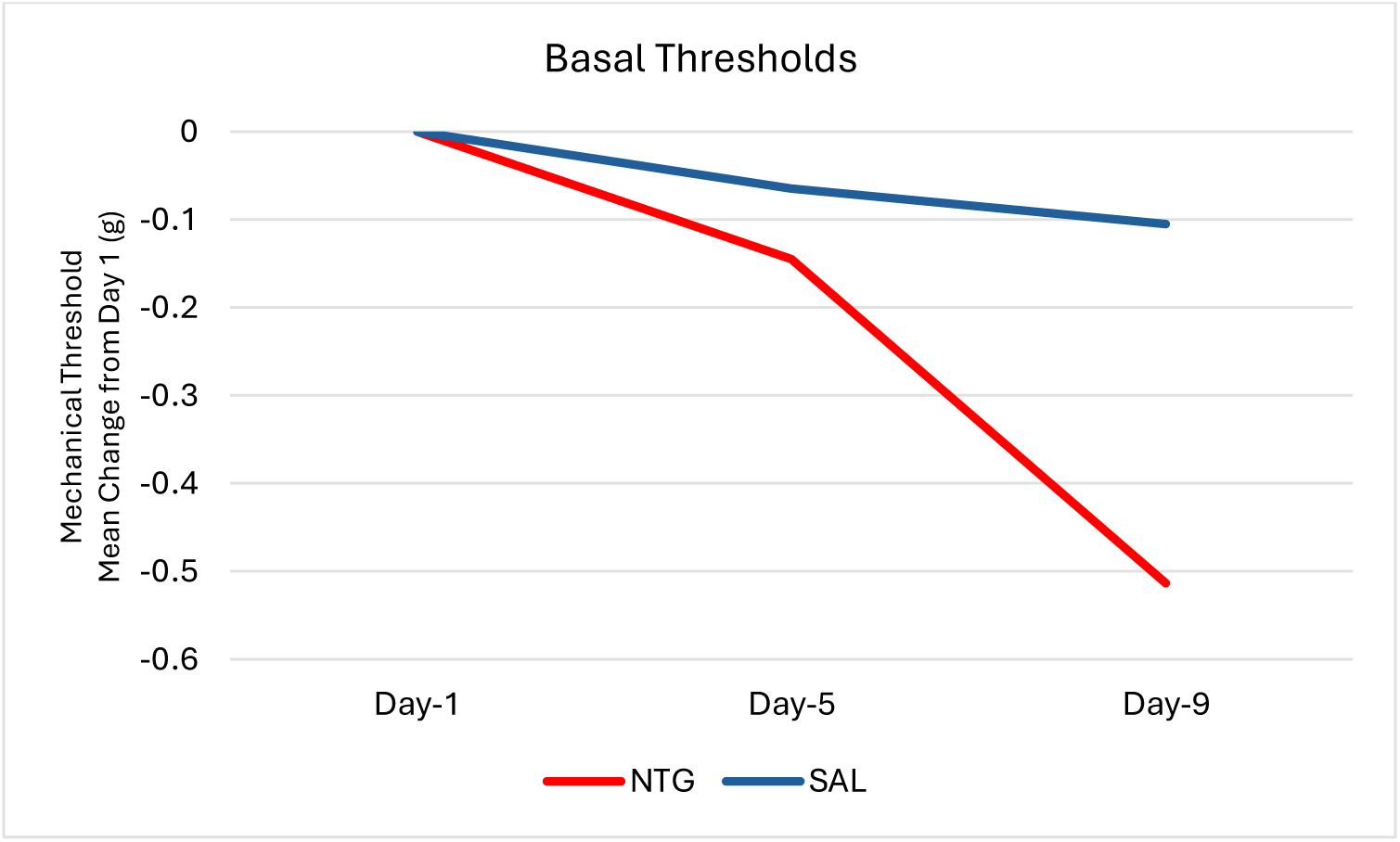
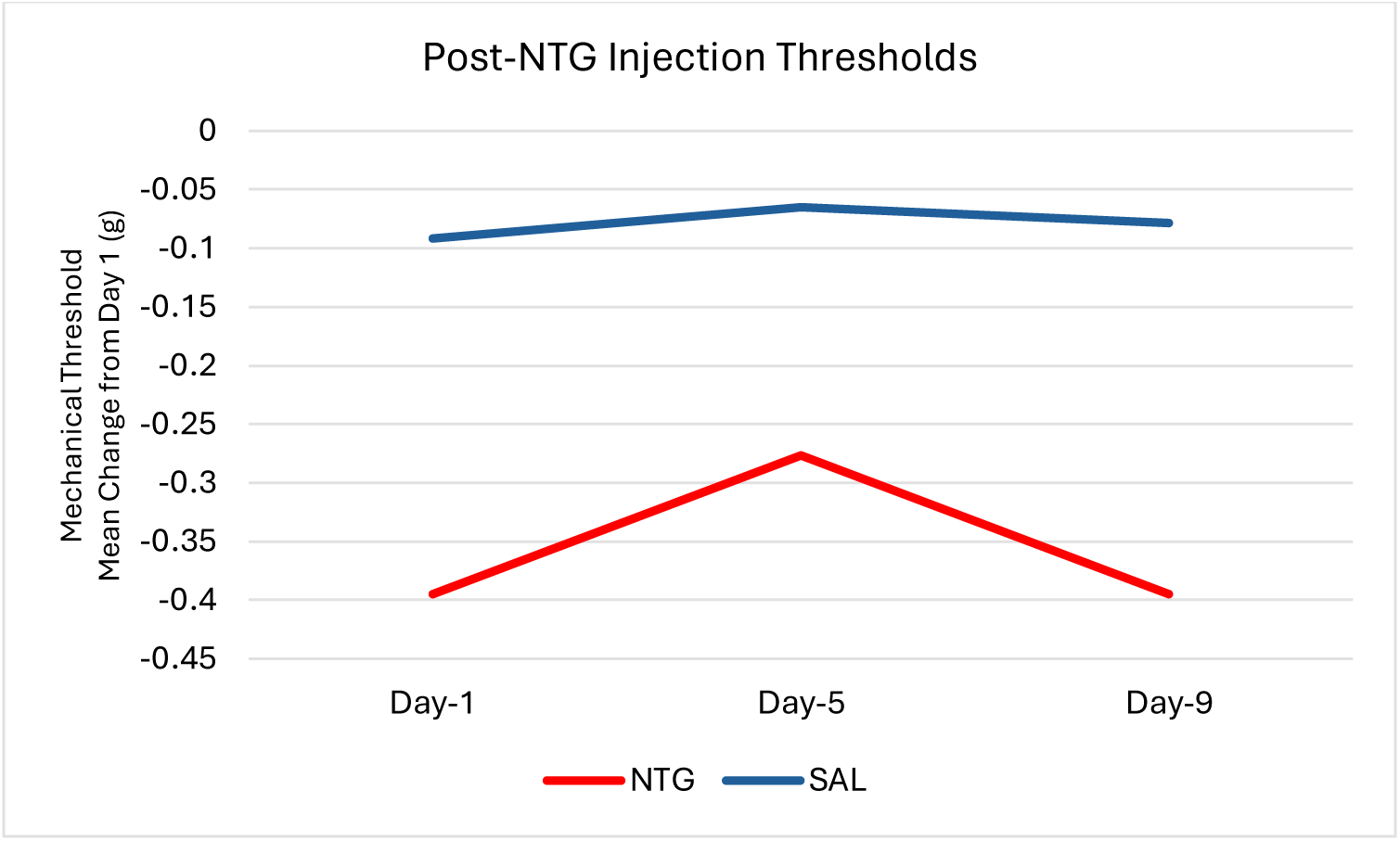
Mechanical Pain Threshold Group Comparison. 1A. Basal Pain Thresholds: Mechanical pain thresholds decreased progressively over time in the NTG group, compared to the SAL group. 1B. NTG-induced Pain Thresholds: Chronic administration of nitroglycerin induces a long-lasting hypersensitivity with reduced mechanical pain thresholds in the NTG group compared to the SAL group. Each timepoint is shown as the average of the mechanical pain thresholds for the six rats in each group. NTG = Nitroglycerine-injected group; Sal = Saline-injected group.

### RNA Sequencing, Circadian Gene Expression, Gene Enrichment Results (Table 1)

1. We analyzed the expression of 13,902 genes in the migraine rat models and identified significant sex-specific differences. Females exhibited a higher number of upregulated genes (338) and fewer downregulated genes (92) compared to males (89 upregulated and 356 downregulated). The top upregulated genes in females included metallothionein-2A (*MT2A,* FC = 36) and solute carrier family 7 (*SLC7A11,* FC = 31) while males displayed robust upregulation of zinc finger and BTB domain containing 16 (*ZBTB16,* FC = 9.6) and S100 calcium binding protein A8 (*S100A8,* FC = 6.23).
2. Conversely, the most downregulated genes in females were High Mobility Group Nucleosome Binding Domain 5 (*HMGN5*, FC = −181), and leukocyte cell derived chemotaxin 1 (*LECT1,* FC = −17) whereas males showed significant downregulation of Secretogranin II (*SCG2*, FC = −13) and glutamate receptor; ionotropic; AMPA 1 (*GRIA1*, FC = −297). Notably, an inverse pattern of gene regulation was observed between sexes, with SLC7A11, SCG2 and GRIA1 being highly upregulated in females and significantly downregulated in males, and highlighting striking sex-specific differences in gene expression.
3. A total of 308 circadian rhythm-associated genes were identified on genecards.org. Of these, 223 (Supplementary Table 1) corresponded to the 13,902 genes expressed in our choroid plexus transcriptomics. The following results related to circadian rhythm were derived from these 223 genes.
4. The following 10 circadian rhythm-associated genes were significantly upregulated in the NTG group compared to the Saline group. Serpin Family E Member 1 (SERPINE1), Mitogen-Activated Protein Kinase 9 (MAPK9), Activating Transcription Factor 4 (ATF4), Lysine Demethylase 5A (KDM5A), G Protein-Coupled Receptor 157 (GPR157), Kruppel-Like Factor 15 (KLF15), Splicing Factor 3A Subunit 3 (SF3A3), Rho-Associated Coiled-Coil Containing Protein Kinase 2 (ROCK2), VAMP-Associated Protein A (VAPA), F-Box And Leucine-Rich Repeat Protein 12 (FBXL12) (Figure 2).

**Figure 2.**
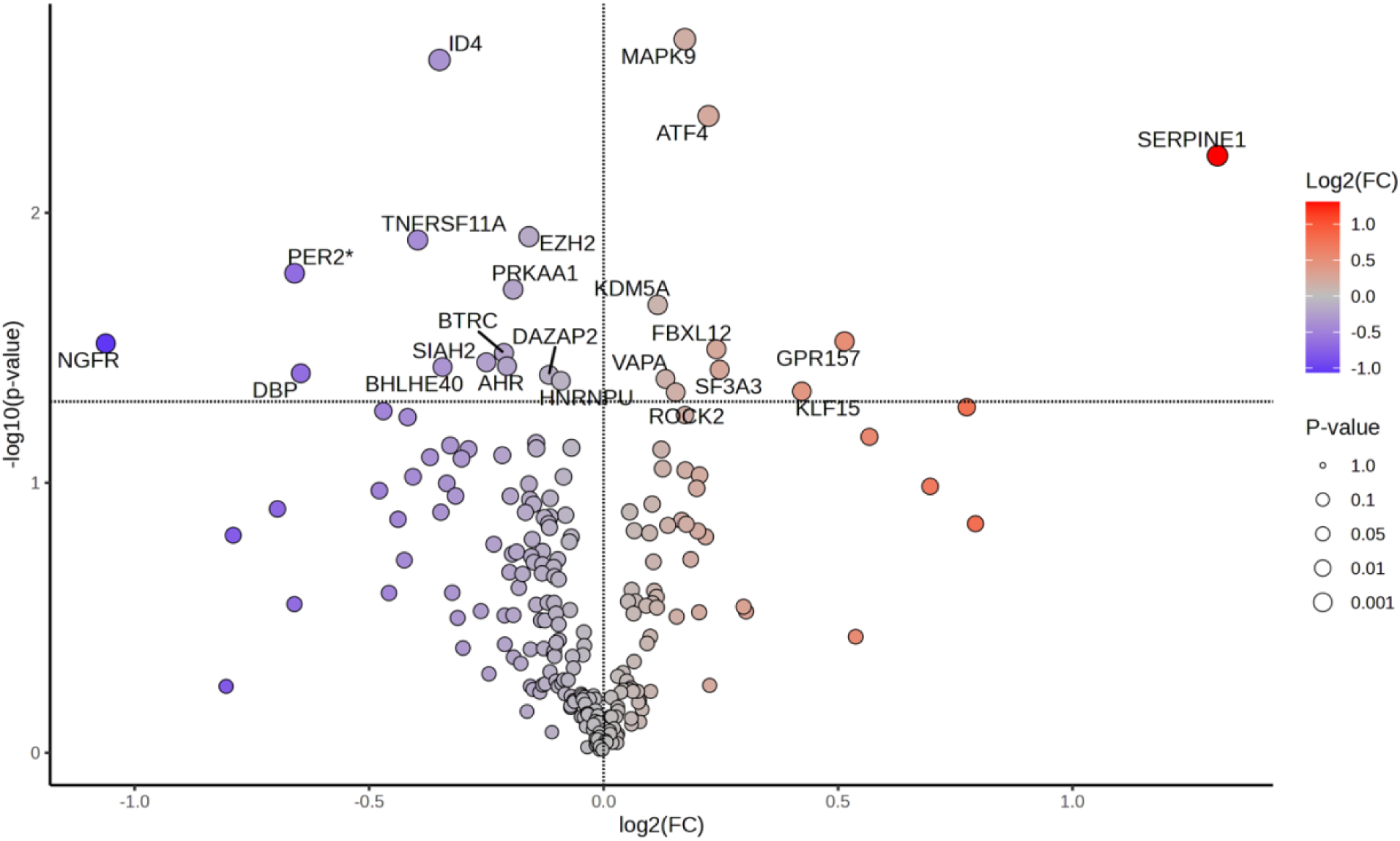
Volcano Plot: Differential Expression of Circadian Rhythm-Associated Genes in NTG vs Saline. x-axis: log2 fold change (log2FC), y-axis: −log10(p-value), with significance threshold at −log10(0.05) = 1.3. Red dots: Upregulated genes (log2FC > 0, FC > 1, p-value < 0.05). Blue dots: Downregulated genes (log2FC < 0, FC < 1, p-value < 0.05). Highlighted genes: Circadian rhythm-associated genes with significant differential expression in the NTG group. Circadian clock genes are marked with an asterisk.
5. The following 13 circadian rhythm-associated genes were significantly downregulated in the NTG group compared to the Saline group. Period Circadian Regulator 2 (PER2)*, Nerve Growth Factor Receptor (NGFR), D-site Albumin Promoter Binding Protein (DBP), Inhibitor of DNA binding 4 (ID4), TNF Receptor Superfamily Member 11a (TNFRSF), Enhancer of Zeste Homolog 2 (EZH2), Protein Kinase AMP-Activated Catalytic Subunit Alpha 1 (PRKAA1), Beta-Transducin Repeat Containing (BTRC), Seven in absentia homolog 2 (SIAH2), Basic Helix-Loop-Helix Family Member E40 (BHLHE40), Aryl Hydrocarbon Receptor (AHR), Heterogeneous Nuclear Ribonucleoprotein U (HNRNPU), DAZ Associated Protein 2 (DAZAP2) (Figure 2). Circadian clock genes are marked with an asterisk.
6. An agglomerative clustering heatmap featuring the 25 most differentially expressed genes successfully distinguished the NTG group from the Saline group with complete accuracy (Figure 3).

**Figure 3.**
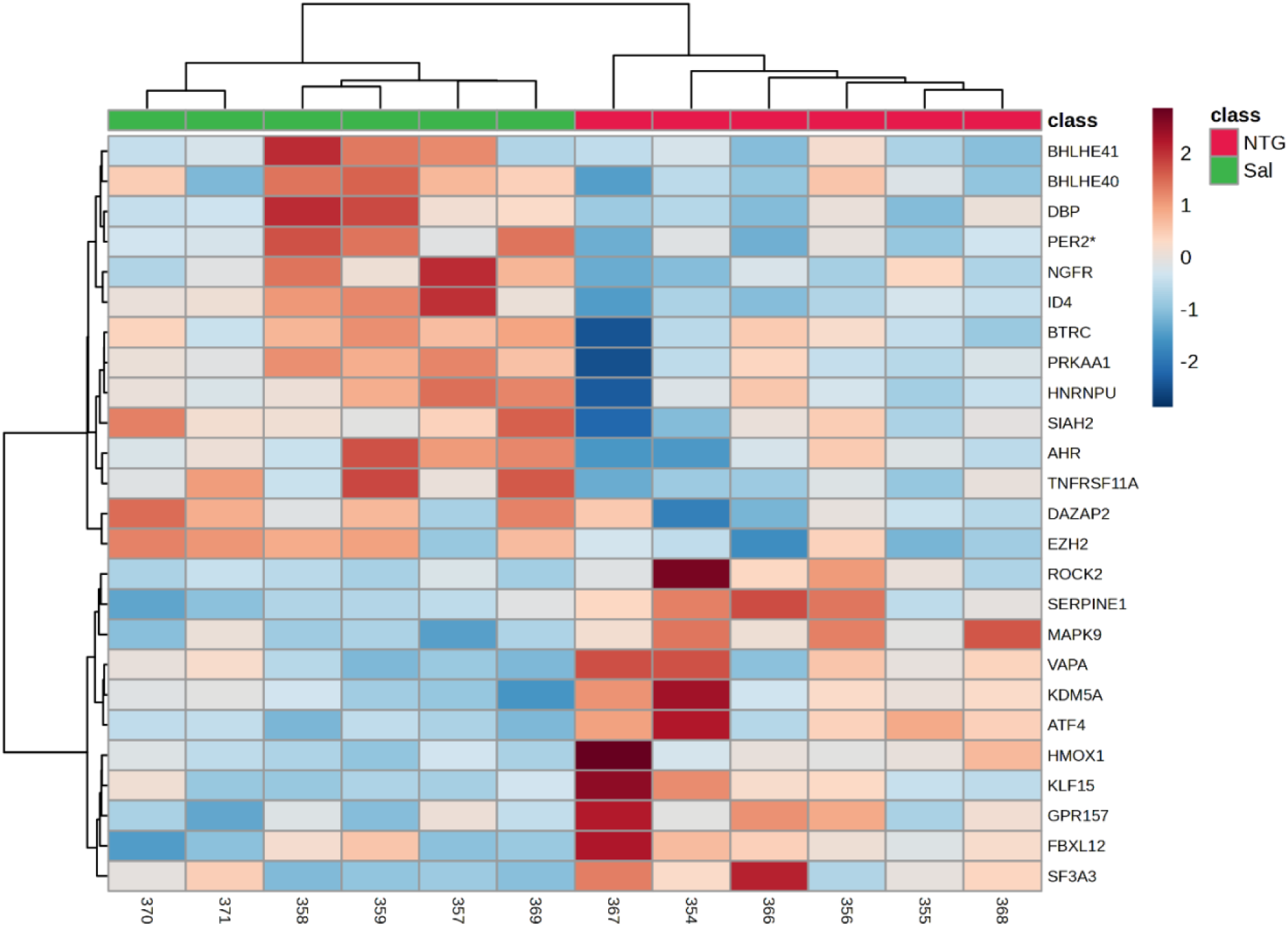
Heatmap: Top 25 Differentially Expressed Circadian Rhythm-Associated Genes in NTG vs. Saline. Rows: Genes (top 25 differentially expressed). Columns: Samples (NTG in red and Saline in green groups). Color scale: log2 expression values (red: high expression, blue: low expression). Clustering: Agglomerative hierarchical clustering of genes and samples based on Euclidean distance and Ward’s linkage method. NTG: Nitroglycerin-injected group. Sal: Saline-treated control group. Circadian clock genes are marked with an asterisk.
7. The following two circadian rhythm-related genes were significantly upregulated in the female NTG group compared to the female saline group. F-Box and Leucine-Rich Repeat Protein 12 (FBXL12) and G Protein-Coupled Receptor 157 (GPR157) (Figure 4).

**Figure 4.**
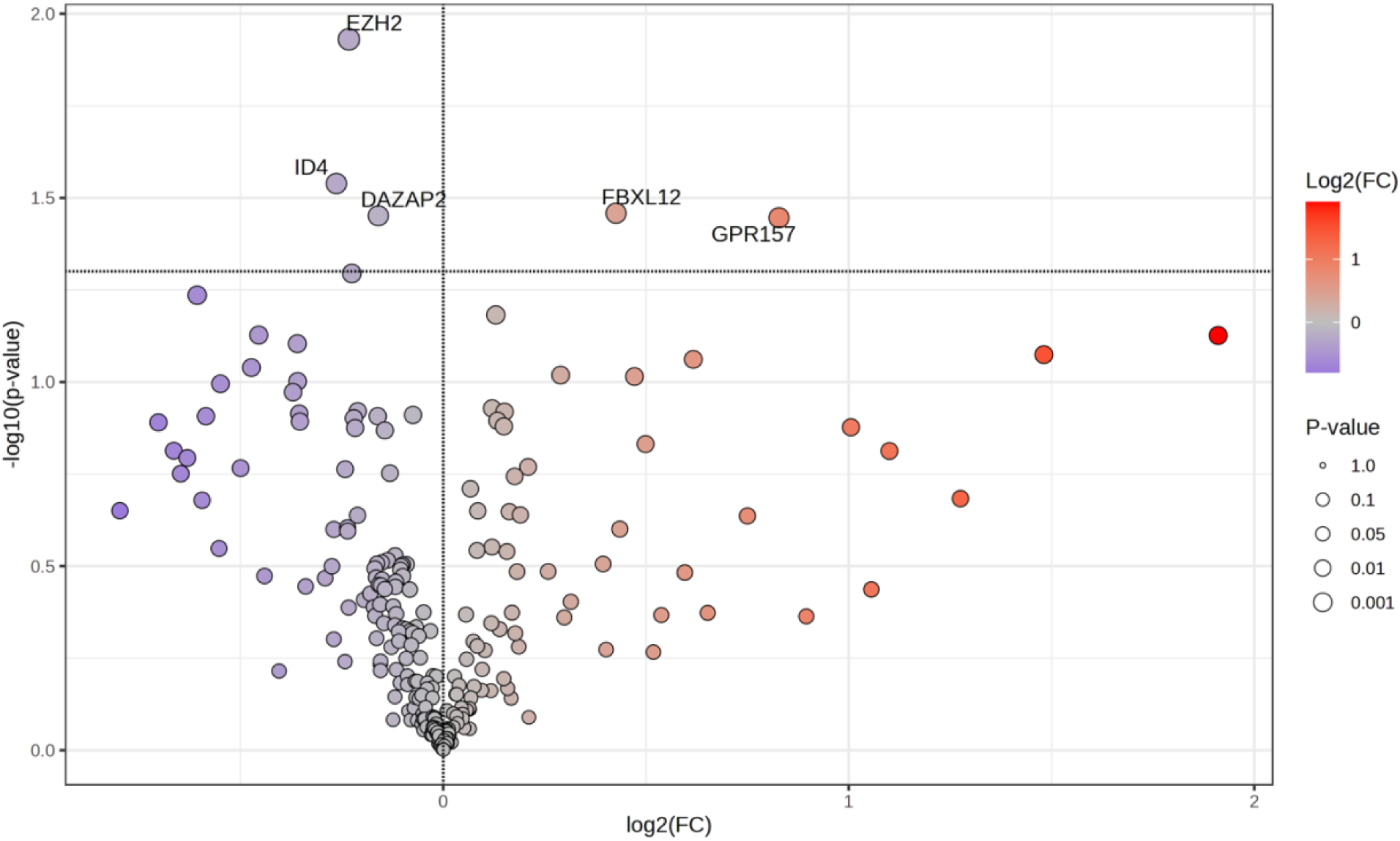
Volcano Plot: Differential Expression of Circadian Rhythm-Associated Genes in Female NTG vs. Female Saline. x-axis: log2 fold change (log2FC), y-axis: −log10(p-value), with significance threshold at −log10(0.05) = 1.3. Red dots: Upregulated genes (log2FC > 0, FC > 1, p-value < 0.05). Blue dots: Downregulated genes (log2FC < 0, FC < 1, p-value < 0.05). Highlighted genes: Circadian rhythm-associated genes with significant differential expression in the NTG group.
8. The following three circadian rhythm-associated genes were significantly downregulated in the female NTG group compared to the female saline group. Enhancer Of Zeste 2 Polycomb Repressive Complex 2 Subunit (EZH2), Inhibitor Of DNA Binding 4 (ID4), DAZ Associated Protein 2 (DAZAP2) (Figure 4).
9. An agglomerative clustering heatmap featuring the 25 most differentially expressed genes distinguished the female NTG group from the female Saline group with complete accuracy (Figure 5).

**Figure 5.**
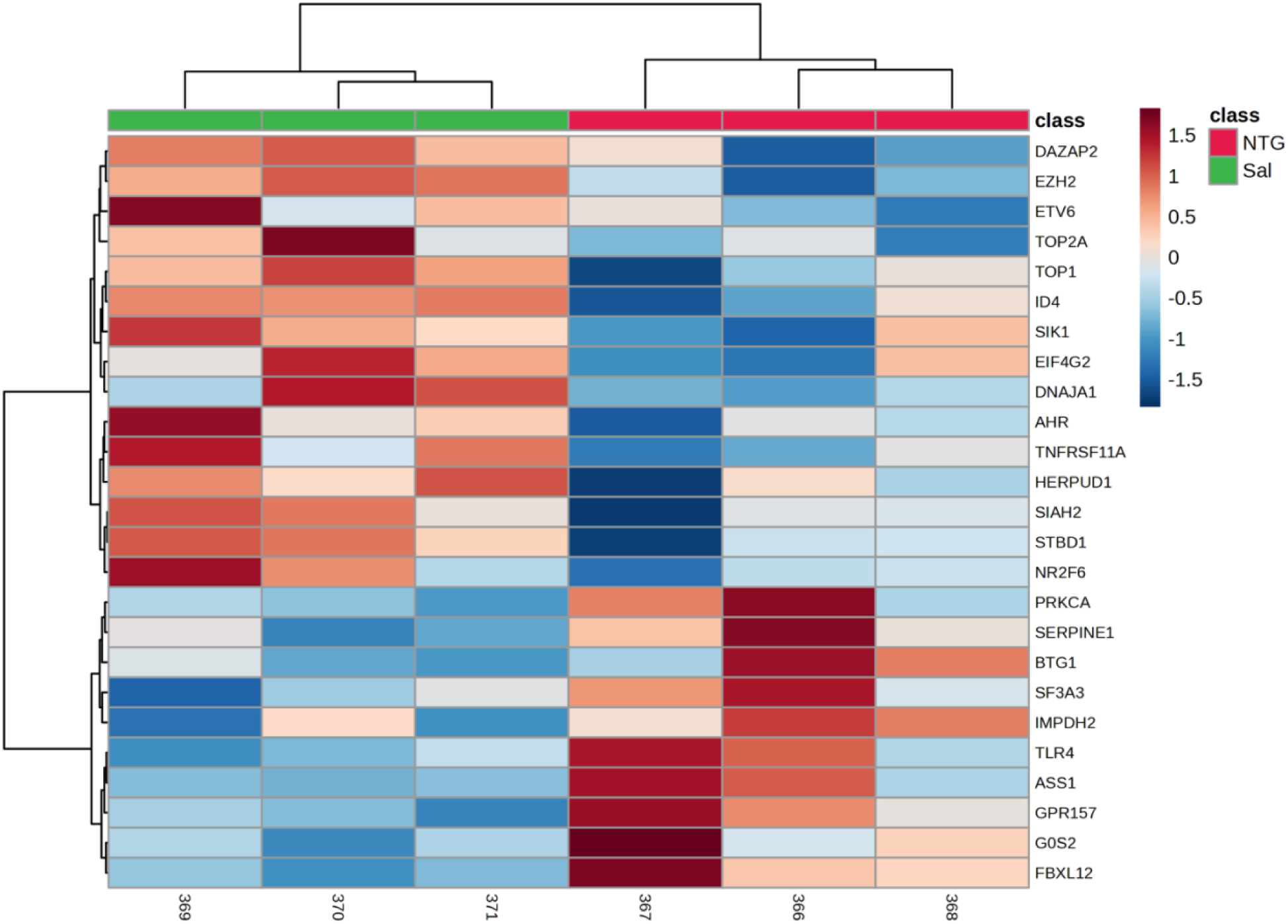
Heatmap: Top 25 Differentially Expressed Circadian Rhythm-Associated Genes in female NTG vs. female Saline. Rows: Genes (top 25 differentially expressed). Columns: Samples (NTG in red and Saline in green groups). Color scale: log2 expression values (red: high expression, blue: low expression). Clustering: Agglomerative hierarchical clustering of genes and samples based on Euclidean distance and Ward’s linkage method. NTG: Nitroglycerin-injected group. Sal: Saline-treated control group.
10. The following 13 circadian rhythm-related genes were significantly upregulated in the male NTG group compared to the male saline group. NK2 Homeobox 1 (NKX2-1), Thymidylate Synthase (TYMS), Protein Phosphatase 1 Catalytic Subunit Gamma (PPP1CC), WD Repeat-Containing Protein 5 (WDR5), Thymidylate Synthase (TYMS), Activating Transcription Factor 4 (ATF4), Trimethylguanosine Synthase 1 (TGS1), Mitogen-Activated Protein Kinase 9 (MAPK9), Circadian Locomotor Output Cycles Kaput (CLOCK*), Ubiquitin A-52 Residue Protein (UBA52), Vesicle-Associated Membrane Protein A (VAPA), Splicing Factor 3A Subunit 3 (SF3A3), and NADH:Ubiquinone Oxidoreductase Subunit A9 (NDUFA9) (Figure 6). Circadian clock genes are marked with an asterisk.
11. The following 22 circadian rhythm-related genes were significantly downregulated in the male NTG group compared to the male saline group. Adenosine Receptor A1 (ADORA1), Cryptochrome 1 (CRY1*), RAR-Related Orphan Receptor B (RORB), Thyroid Hormone Receptor Associated Protein 3 (THRAP3), NCK Associated Protein 1 (NCKAP1), Beta-Transducin Repeat Containing (BTRC), Protein Kinase AMP-Activated Alpha 1 (PRKAA1), Chromodomain Helicase DNA Binding Protein 9 (CHD9), SIN3 Transcription Regulator Family Member A (SIN3A), Casein Kinase 1 Epsilon (CSNK1E*), Helicase With Zinc Finger 2 (HELZ2), Nuclear Receptor Coactivator 1 (NCOA1), Zinc Finger Homeobox 3 (ZFHX3), Nuclear Receptor Coactivator 2 (NCOA2), Nuclear Receptor Subfamily 1 Group H Member 3 (NR1H3), Basic Helix-Loop-Helix Family Member E40 (BHLHE40), Carnitine Palmitoyltransferase 1A (CPT1A), PH Domain Leucine-Rich Repeat Protein Phosphatase 1 (PHLPP1), Protein Kinase C Alpha (PRKCA), Inhibitor Of DNA Binding 4 (ID4), Myocyte Enhancer Factor 2C (MEF2C), and Basic Helix-Loop-Helix Family Member E41 (BHLHE41) (Figure 6). Circadian clock genes are marked with an asterisk.

**Figure 6.**
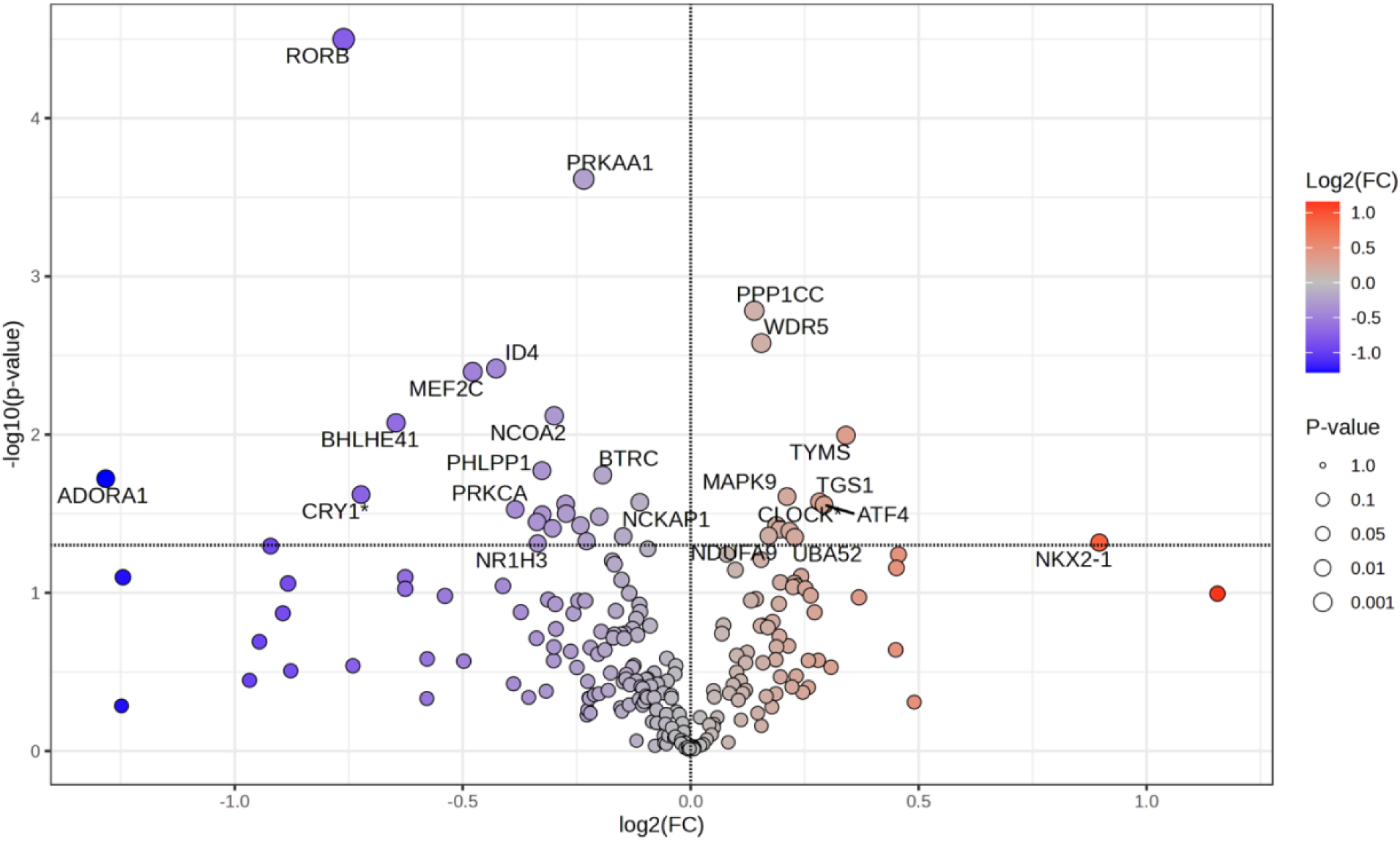
Volcano Plot: Differential Expression of Circadian Rhythm-Associated Genes in Male NTG vs. Male Saline. x-axis: log2 fold change (log2FC), y-axis: −log10(p-value), with significance threshold at −log10(0.05) = 1.3. Red dots: Upregulated genes (log2FC > 0, FC > 1, p-value < 0.05). Blue dots: Downregulated genes (log2FC < 0, FC < 1, p-value < 0.05). Highlighted genes: Circadian rhythm-associated genes with significant differential expression in the NTG group. Circadian clock genes are marked with an asterisk.
12. An agglomerative clustering heatmap featuring the 25 most differentially expressed genes distinguished the male NTG group from the male Saline group with complete accuracy (Figure 7).

**Figure 7.**
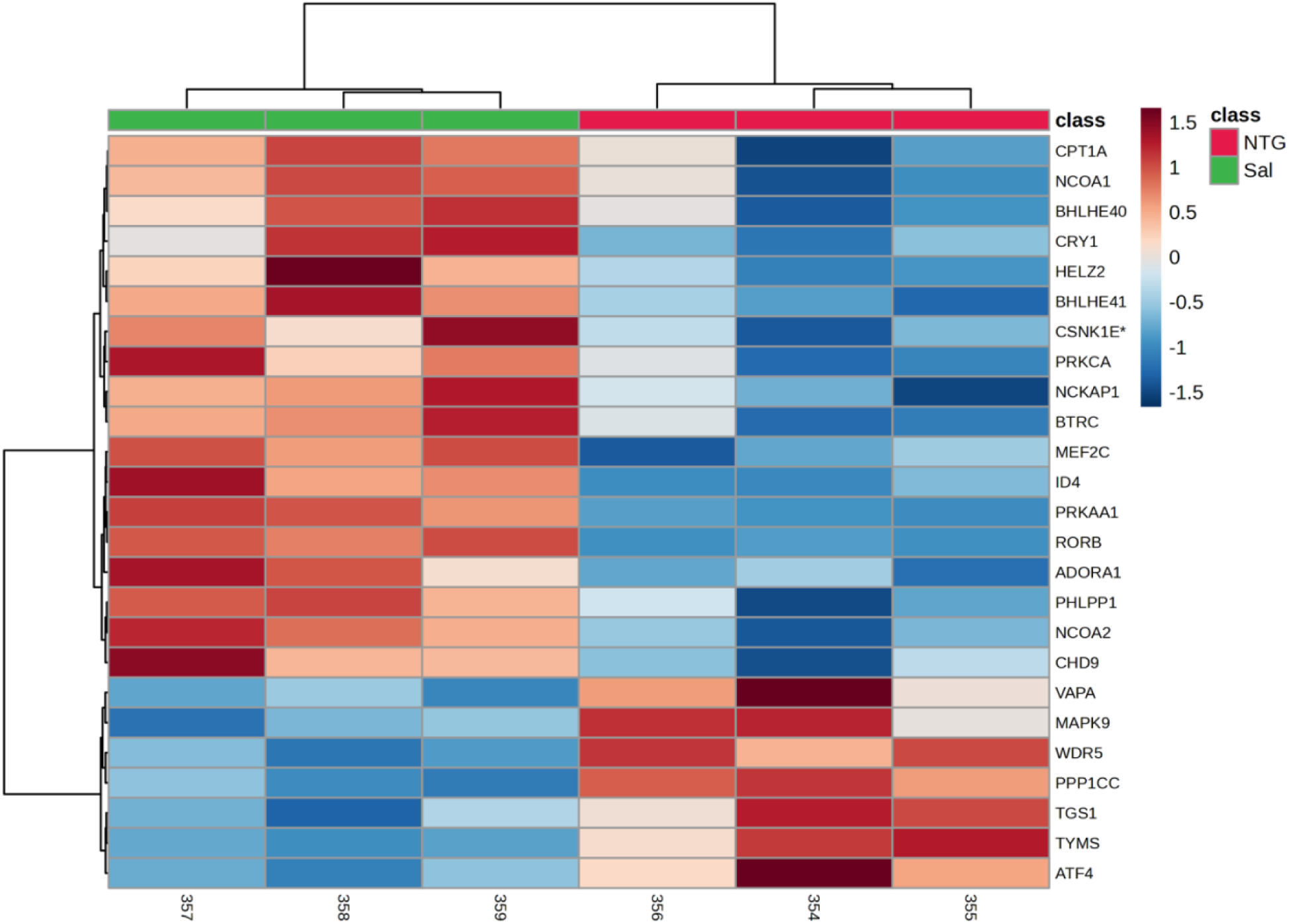
Heatmap: Top 25 Differentially Expressed Circadian Rhythm-Associated Genes in Male NTG vs. Male Saline. Rows: Genes (top 25 differentially expressed). Columns: Samples (NTG in red and Saline in green groups). Color scale: log2 expression values (red: high expression, blue: low expression). Clustering: Agglomerative hierarchical clustering of genes and samples based on Euclidean distance and Ward’s linkage method. NTG: Nitroglycerin-injected group. Sal: Saline-treated control group.
13. GO enrichment analysis of Biological Processes revealed that the top 3 pathways enriched among upregulated genes in the NTG vs SAL comparison were related to circadian rhythm, specifically: regulation of circadian rhythm, circadian regulation of gene expression, and circadian rhythm. GO enrichment analysis revealed that the top enriched Cellular Components were the SCF ubiquitin ligase complex and the RNA polymerase II transcription regulator complex. Furthermore, the Molecular Function of GO analysis showed significant enrichment of terms related to transcription coregulator binding and RNA polymerase II-specific DNA-binding transcription factor binding (Supplementary Figure 1A-C).
14. KEGG enrichment analysis revealed that the top four enriched pathways in the NGT vs. SAL comparison were those involved in circadian rhythm regulation, insulin resistance, glucagon signaling, and amphetamine addiction. Reactome analysis revealed that regulation of lipid metabolism by PPARα and heme signaling was the top two pathways. (Supplementary Figure 1D-E).
15. KEGG enrichment in females revealed the HIF-1 (Hypoxia-Inducible Factor signaling) pathway. GO Biological Process in females revealed regulation of circadian and wound healing, involved in inflammatory response as the top two pathways. GO Molecular Function in females showed haptoglobin binding and hemoglobin alpha binding as the top two pathways. GO Cellular Component in females showed hemoglobin complex and haptoglobin-hemoglobin complex as the top two pathways (Supplementary Figure 2A-D).
16. KEGG enrichment in males revealed the circadian rhythm pathway. GO Biological Process in males revealed arachidonic acid metabolite production and leukotriene production, both involved in inflammatory response, as the top two pathways. GO Molecular Function in males showed heterotrimeric G-protein binding and hemi-methylated DNA-binding as the top two pathways. GO Cellular Component in males showed no overrepresented pathways (Supplementary Figure 3A-C).
17. Reactome pathway analysis yielded no significant pathway enrichment for the DEGs in both males and females, likely due to the limited number of DEGs and small sample size.

## Discussion

RNA sequencing of 13,902 genes in choroid plexus tissue from the 4^th^ ventricle of nitroglycerin (NTG)-induced migraine rat models unveiled profound sex-specific differences in gene expression, revealing distinct molecular landscapes intricately woven with circadian rhythm dysregulation—a hallmark of migraine pathology. The choroid plexus, a pivotal regulator of cerebrospinal fluid (CSF) production, maintains brain homeostasis through a robust circadian clock that orchestrates diurnal fluctuations in CSF composition and intracranial pressure, both implicated in migraine’s throbbing pain and sensory disturbances. The choroid plexus controls the influx of molecules and inflammatory mediators into the brain through the blood-cerebrospinal fluid barrier (BCB). Unlike the blood-brain barrier (BBB), the BCB is characterized by fenestrated capillaries and gap junctions, facilitating the transport of larger macromolecules and immune cells.

**Table 1:** Differentially Expressed Genes in Choroid Plexus, Including Circadian Rhythm-Associated Genes, in NTG vs. Saline and by Sex.

| Genes | Function | Differential Expression (NTG vs. Saline) | Female Change | Male Change |
| --- | --- | --- | --- | --- |
| *MT2A*,<br>*SLC7A11* | Antioxidant | ↑ | ↑ | ↔<br>(*SLC7A11* ↓) |
| *ZBTB16*,<br>*S100A8* | Anti-inflammatory,<br>transcriptional factors | ↑ | ↔ | ↑ |
| *SERPINE1*,<br>*ATF4* | Circadian rhythm | ↑ | ↑ | ↑ |
| *PER2*, *CRY1*,<br>*DBP* | Circadian rhythm | ↓ | ↓ | ↓ |
| *CLOCK* | Circadian rhythm | ↑ | ↔ | ↑ |
| *FBXL12*,<br>*GPR157* | Circadian rhythm | ↑ | ↑ | ↔ |
| *NKX2-1*,<br>*CSNK1E* | Circadian rhythm | ↑ | ↔ | ↑ |

Females exhibited a striking bias toward upregulation (338 genes increased, 92 downregulated), contrasting sharply with males (89 upregulated, 356 downregulated), suggesting sex-dependent choroid plexus responses—females favoring CSF-mediated neuroprotection and males amplifying inflammation or suppression—tightly coupled to circadian timing. Among 308 circadian-associated genes identified via GeneCards, 223 overlapped with our choroid plexus transcriptomics, and agglomerative clustering heatmaps of the top 25 differentially expressed genes unequivocally distinguished NTG from saline groups across both sexes (Figures 3, 5, 7). Functional enrichment analyses cemented circadian rhythm regulation, transcriptional control, and metabolic pathways as foundational to migraine’s chronobiological underpinnings, positioning the 4^th^ ventricle choroid plexus as a critical circadian nexus in brain pathology.

### Sex-Specific Gene Regulation

In female 4^th^ ventricle choroid plexus tissue, upregulated genes illuminated a circadian-orchestrated protective fortress against migraine triggers. Metallothionein-2A (*MT2A*, FC = 36) encodes a cysteine-rich protein that sequesters zinc and scavenges reactive oxygen species (ROS), countering oxidative stress—a migraine catalyst peaking diurnally under *CLOCK/BMAL1* regulation—potentially stabilizing CSF redox balance to shield brainstem and cerebellar regions proximal to the 4^th^ ventricle. Solute carrier family 7 member 11 (*SLC7A11*, FC = 31) imports cystine for glutathione synthesis via the cystine/glutamate antiporter, fortifying CSF antioxidant defenses against glutamate excitotoxicity in a rhythm synchronized with diurnal redox oscillations (Figure 2). Downregulation was restrained, with high mobility group nucleosome binding domain 5 (*HMGN5*, FC = −181) reducing chromatin accessibility—a process governed by circadian remodeling cycles in choroid plexus epithelial cells—and leukocyte cell-derived chemotaxin 1 (*LECT1*, FC = −17) suppressing angiogenesis, subtly adjusting CSF dynamics through diurnal vascular rhythms.

Males, conversely, showcased a divergent molecular signature in choroid plexus tissue, with upregulated genes driving transcriptional and inflammatory cascades. Zinc finger and BTB domain containing 16 (ZBTB16) encodes a transcription factor modulating differentiation and stress responses, likely interacting with circadian repressors like PER2 to rewire CSF gene expression, while S100 calcium binding protein A8 (S100A8) forms calprotectin with S100A9, amplifying inflammation via TLR4 signaling—aligned with diurnal immune patterns in choroid plexus macrophages—suggesting a male-specific inflammatory surge in CSF composition (Figure 6). Downregulation was stark, with secretogranin II (SCG2, FC = −13) crippling neuropeptide signaling (e.g., secretoneurin) essential for circadian pain modulation in CSF, and glutamate receptor, ionotropic, AMPA 1 (GRIA1, FC = −297) diminishing excitatory synaptic transmission, tied to diurnal glutamate cycles influencing brainstem signaling via CSF.

### Circadian Rhythm Alterations

Circadian gene expression in 4^th^ ventricle choroid plexus tissue was profoundly disrupted in NTG versus saline controls, with 10 genes upregulated and 13 downregulated among the 223 circadian-associated genes (Figure 2). The choroid plexus clock, a master regulator of CSF secretion, drives diurnal variations in intracranial pressure and solute transport—key migraine modulators. Upregulated genes signaled a shift in this temporal orchestration: serpin family E member 1 (SERPINE1) peaks nocturnally, exacerbating CSF vascular dysfunction; mitogen-activated protein kinase 9 (MAPK9) fuels inflammation via JNK signaling with CLOCK interactions; and activating transcription factor 4 (ATF4) governs stress responses in diurnal cycles. Lysine demethylase 5A (KDM5A) represses genes epigenetically, G protein-coupled receptor 157 (GPR157) alters sensory signaling, kruppel-like factor 15 (KLF15) tunes metabolism as a clock output, splicing factor 3A subunit 3 (SF3A3) enhances RNA processing, rho-associated coiled-coil containing protein kinase 2 (ROCK2) regulates vascular tone, VAMP-associated protein A (VAPA) supports vesicle trafficking, and F-box and leucine-rich repeat protein 12 (FBXL12) degrades clock proteins—all disrupting CSF rhythmicity (Figure 3 heatmap).

Downregulated genes underscored a collapse of choroid plexus clock integrity: period circadian regulator 2 (PER2*, asterisked) represses CLOCK/BMAL1, and its loss dismantles CSF rhythm amplitude; nerve growth factor receptor (NGFR) sustains neuronal survival diurnally; D-site albumin promoter binding protein (DBP) drives metabolic oscillations; and inhibitor of DNA binding 4 (ID4) curbs transcription. TNF receptor superfamily member 11A (TNFRSF11A) modulates inflammation, enhancer of zeste homolog 2 (EZH2) silences genes epigenetically, protein kinase AMP-activated catalytic subunit alpha 1 (PRKAA1) regulates energy, beta-transducin repeat containing (BTRC) targets clock proteins, seven in absentia homolog 2 (SIAH2) ubiquitinates substrates, basic helix-loop-helix family member E40 (BHLHE40) represses clock genes, aryl hydrocarbon receptor (AHR) links environmental cues, heterogeneous nuclear ribonucleoprotein U (HNRNPU) processes RNA, and DAZ-associated protein 2 (DAZAP2) supports splicing—all with circadian roles in CSF dynamics.

### Sex-Specific Circadian Gene Regulation

In female NTG choroid plexus tissue, FBXL12 and GPR157 were upregulated (Figure 4), destabilizing CSF rhythms and shifting sensory signaling, while EZH2, ID4, and DAZAP2 were downregulated, easing epigenetic and splicing control (Figure 5 heatmap). In males, 13 genes were upregulated (Figure 6), including:

- NK2 homeobox 1 (NKX2-1) and circadian locomotor output cycles kaput (CLOCK)*: Amplify CSF clock output.
- Thymidylate synthase (TYMS) and trimethylguanosine synthase 1 (TGS1): Bolster DNA/RNA synthesis.
- Protein phosphatase 1 catalytic subunit gamma (PPP1CC), WD repeat-containing protein 5 (WDR5), ubiquitin A-52 residue protein (UBA52), and NADH:ubiquinone oxidoreductase subunit A9 (NDUFA9): Modulate cellular processes.
- ATF4, MAPK9, VAPA, and SF3A3: Enhance stress and RNA processing.

Males downregulated 22 genes (Figure 6), including adenosine receptor A1 (ADORA1), cryptochrome 1 (CRY1*), casein kinase 1 epsilon (CSNK1E*), RAR-related orphan receptor B (RORB), thyroid hormone receptor-associated protein 3 (THRAP3), NCK-associated protein 1 (NCKAP1), BTRC, PRKAA1, chromodomain helicase DNA binding protein 9 (CHD9), SIN3 transcription regulator family member A (SIN3A), helicase with zinc finger 2 (HELZ2), nuclear receptor coactivator 1 (NCOA1), zinc finger homeobox 3 (ZFHX3), nuclear receptor coactivator 2 (NCOA2), nuclear receptor subfamily 1 group H member 3 (NR1H3), BHLHE40, carnitine palmitoyltransferase 1A (CPT1A), PH domain leucine-rich repeat protein phosphatase 1 (PHLPP1), protein kinase C alpha (PRKCA), ID4, myocyte enhancer factor 2C (MEF2C), and basic helix-loop-helix family member E41 (BHLHE41), signaling a profound CSF rhythm collapse (Figure 7 heatmap).

### Overlaps and Inverse Regulation

A rigorous analysis of overlaps (same direction) and inverse regulations (opposite direction) in 4th ventricle choroid plexus tissue reveals critical sex-specific patterns among all differentially expressed and circadian-associated genes:

#### Overlaps (Same Direction)

- General Genes: No top differentially expressed genes (MT2A, SLC7A11, HMGN5, LECT1 in females; ZBTB16, S100A8, SCG2, GRIA1 in males) overlap in the same direction, highlighting distinct choroid plexus responses.
- Circadian Genes:

- ATF4: Upregulated in NTG vs. saline and males, amplifying stress responses in CSF with diurnal unfolded protein cycles.
- MAPK9: Upregulated in NTG vs. saline and males, driving inflammation via CLOCK in choroid plexus CSF regulation.
- VAPA: Upregulated in NTG vs. saline and males, supporting CSF vesicle trafficking diurnally.
- SF3A3: Upregulated in NTG vs. saline and males, enhancing RNA processing with circadian splicing rhythms.
- FBXL12: Upregulated in NTG vs. saline and females, destabilizing CSF rhythms via clock protein degradation.
- GPR157: Upregulated in NTG vs. saline and females, shifting CSF sensory signaling diurnally.
- ID4: Downregulated in NTG vs. saline, females, and males, relaxing transcriptional repression across sexes, disrupting choroid plexus clock control.
- EZH2: Downregulated in NTG vs. saline and females, easing epigenetic silencing in CSF production.
- DAZAP2: Downregulated in NTG vs. saline and females, impairing RNA splicing in CSF.
- BTRC: Downregulated in NTG vs. saline and males, altering clock protein turnover in CSF.
- PRKAA1: Downregulated in NTG vs. saline and males, disrupting energy homeostasis in CSF.
- BHLHE40: Downregulated in NTG vs. saline and males, weakening clock repression in CSF regulation.

#### Inverse Regulation

- General Genes:

- SLC7A11: Upregulated in females (FC = 31), fortifying CSF antioxidant defenses diurnally, but downregulated in males (implied from inverse note), diminishing protection against excitotoxicity.
- SCG2: Upregulated in females (implied), sustaining CSF neuropeptide signaling for pain modulation, but downregulated in males (FC = −13), crippling diurnal pain control.
- GRIA1: Upregulated in females (implied), boosting CSF-mediated synaptic activity diurnally, but downregulated in males (FC = −297), dampening glutamate signaling.
- Circadian Genes: No explicit inverse regulations exist within the sex-specific circadian lists (female: FBXL12, GPR157 up; EZH2, ID4, DAZAP2 down; male: 13 up, 22 down). For instance:

- FBXL12: Upregulated in females, not downregulated in males.
- CLOCK: Upregulated in males, not downregulated in females.
- ID4: Downregulated in both, not inverse.

Broader NTG vs. saline trends (e.g., PER2 down, CLOCK up) lack sex-specific data to confirm inversions (e.g., CLOCK). Thus, no direct circadian gene inversions are evident, unlike the general genes.

### Pathway Enrichment Insights

GO enrichment analysis of DEGs in the choroid plexus of NTG-treated versus saline-treated rats revealed significant enrichment of circadian rhythm, its regulation, and rhythmic gene expression as the top biological processes (Supplementary Figure 1A), underscoring clock dysregulation as a critical factor in CSF production and migraine pathophysiology. Enriched cellular components, including the SCF ubiquitin ligase complex and RNA polymerase II transcription regulator complex, alongside molecular functions such as transcription coregulator binding and RNA polymerase II-specific DNA-binding, suggest transcriptional and post-translational reprogramming in the NTG model (Supplementary Figure 1B-C). KEGG pathway analysis revealed significant enrichment of circadian rhythm regulation, insulin resistance, glucagon signaling, and amphetamine addiction (Supplementary Figure 1D), linking neurotransmitter dysregulation and metabolic stress to choroid plexus chronobiology. Reactome analysis highlighted regulation of lipid metabolism by PPARα and heme signaling as key pathways in the combined NTG vs. saline comparison (Supplementary Figure 1E), implicating lipid homeostasis and oxygen transport in migraine-related choroid plexus dysfunction. Sex-specific analyses uncovered distinct molecular profiles: females exhibited enrichment of HIF-1 signaling, hemoglobin-related pathways (haptoglobin and hemoglobin alpha binding, hemoglobin complex), and wound healing/inflammatory response processes (Supplementary Figure 2A-D), indicating hypoxia-driven and inflammatory contributions to migraine in females. In contrast, males showed enrichment of arachidonic acid and leukotriene production pathways, both associated with inflammatory responses, with molecular functions such as heterotrimeric G-protein binding and hemi-methylated DNA-binding (Supplementary Figure 3A-C). However, no overrepresented cellular components were identified (Supplementary Figure 3D). Reactome analysis failed to identify significant pathway enrichment in sex-specific DEGs, likely due to the limited number of DEGs and small sample size (n=3; Supplementary Figures 2D, 3D). These findings collectively highlight sex-specific circadian and metabolic disruptions in the choroid plexus, with female-specific hypoxia and hemoglobin pathways and male-specific inflammatory lipid pathways, suggesting novel therapeutic targets for personalized migraine interventions.

### Integration and Implications

The inverse regulation of SLC7A11, SCG2, and GRIA1 in 4th ventricle choroid plexus tissue underscores a sex-specific molecular dichotomy: females harness diurnal antioxidant and synaptic support to fortify CSF, while males suppress these, amplifying inflammation (S100A8) and transcriptional shifts (ZBTB16). Overlaps like ID4 and ATF4 signal shared circadian disruptions and stress responses in CSF regulation; yet, the absence of inverse circadian gene regulation suggests that sex-specific clock alterations (e.g., CLOCK in males, FBXL12 in females) diverge without direct opposition. This choroid plexus chronopathology, validated by heatmaps and enrichment, positions circadian misalignment as a driver of migraine, advocating for sex-tailored chronotherapies—e.g., targeting diurnal CSF antioxidant peaks in females or clock restoration in males—to recalibrate these molecular rhythms with scientific precision.

The sexual dimorphism observed in our study highlights the importance of considering sex differences in the development of therapeutic strategies for chronic migraine. The distinct sets of differentially expressed genes in females and males suggest that different genetic substrates of the molecular mechanisms may contribute to the development of chronic migraine in each sex. These findings have significant implications for the development of sex-specific therapeutic strategies that target the regulation of circadian rhythms in the treatment of chronic migraine.

The glymphatic system, a network in the brain responsible for clearing waste products, is intricately connected to the circadian rhythm. The most effective waste removal occurs during sleep and is primarily controlled by the suprachiasmatic nucleus, the circadian master clock, in response to light-dark cycles. The choroid plexus is the crucial starting point for the glymphatic system, integral to maintaining brain health. In our study, the choroid plexus of chronic migraine models exhibits an overall increased dysregulation of genes encoding proteins that reset the circadian system, specifically circadian clock genes, ribosomal proteins, spermidine, and other proteins related to the circadian rhythm. This dysregulation in non-neuronal cells may act as a mechanism for recurrent migraine attacks, potentially altering glymphatic function in migraine. The sex-specific differences may explain the sex disparity in migraine burden and related mechanisms.

Our study’s findings are limited by the use of a rat model, which may not fully recapitulate the complexity of human chronic migraine. Further studies are needed to validate our findings in human populations and to explore the potential therapeutic implications of targeting circadian rhythm-associated genes in chronic migraine. Single-cell analysis and adjustment for the estrous cycle can enhance the robustness of our preliminary findings.

In conclusion, our study underscores the significance of accounting for sex differences in gene expression and circadian rhythm-associated gene regulation when examining the pathophysiology of chronic migraine. The identification of robust biomarkers and therapeutic targets, as well as the elucidation of the molecular mechanisms underlying chronic migraine, may provide novel avenues for the development of effective therapeutic strategies for this debilitating disorder. Furthermore, our study provides new insights into the molecular mechanisms underlying chronic migraine, revealing distinct sets of differentially expressed genes in females and males.

These findings provide a foundation for the development of sex-specific therapeutic strategies that target the regulation of circadian rhythms in the treatment of chronic migraine.

### Limitation

The primary limitation is the small sample size (3 females, 3 males per group), which may limit statistical power. However, the results provide valuable insights to guide future studies with larger cohorts.

---

**Supplementary Figure 1A.**
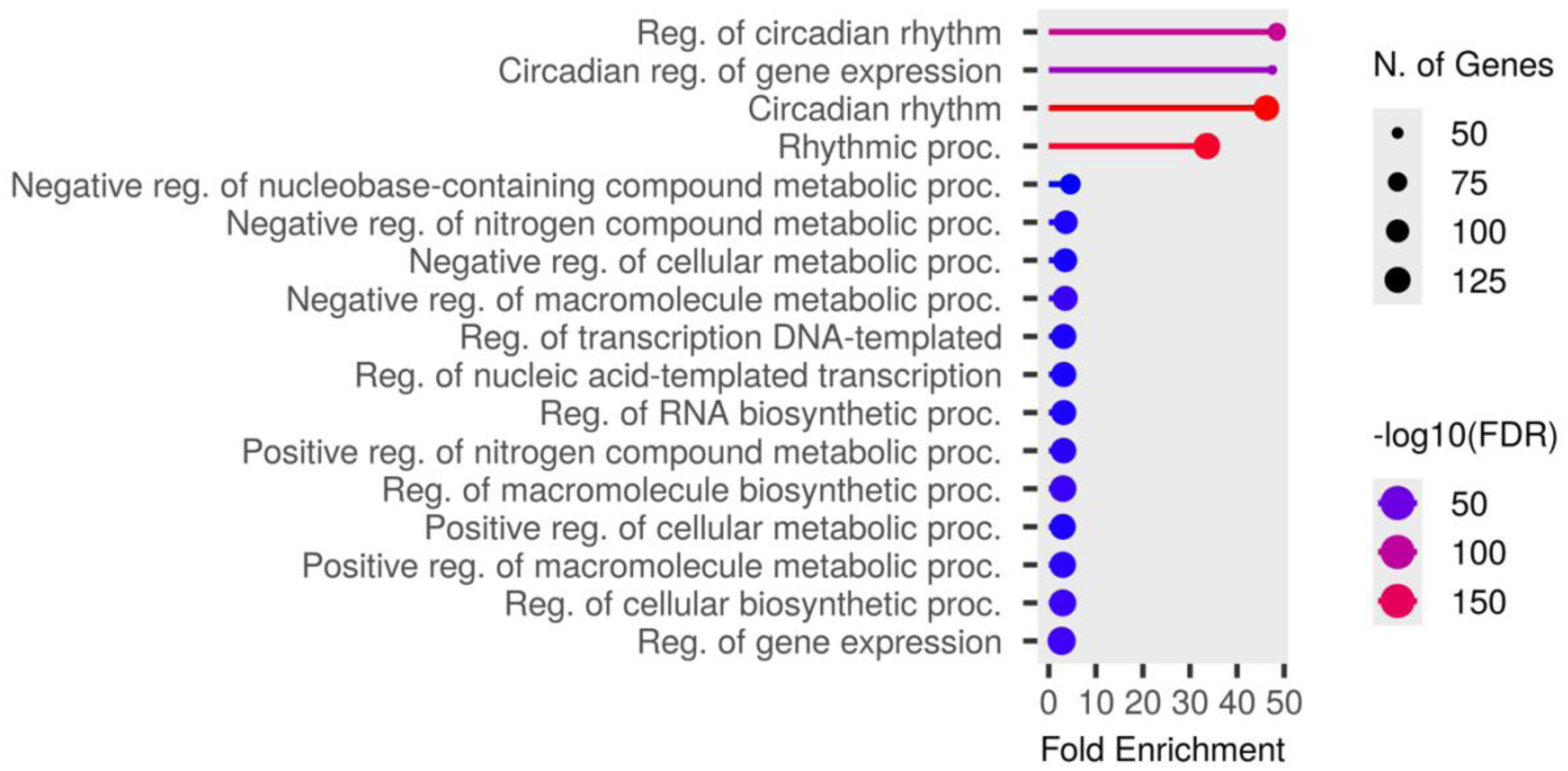
Gene Ontology (GO) Analysis of Biological Processes in NTG-induced rat migraine models vs. saline-injected controls.

**Supplementary Figure 1B.**
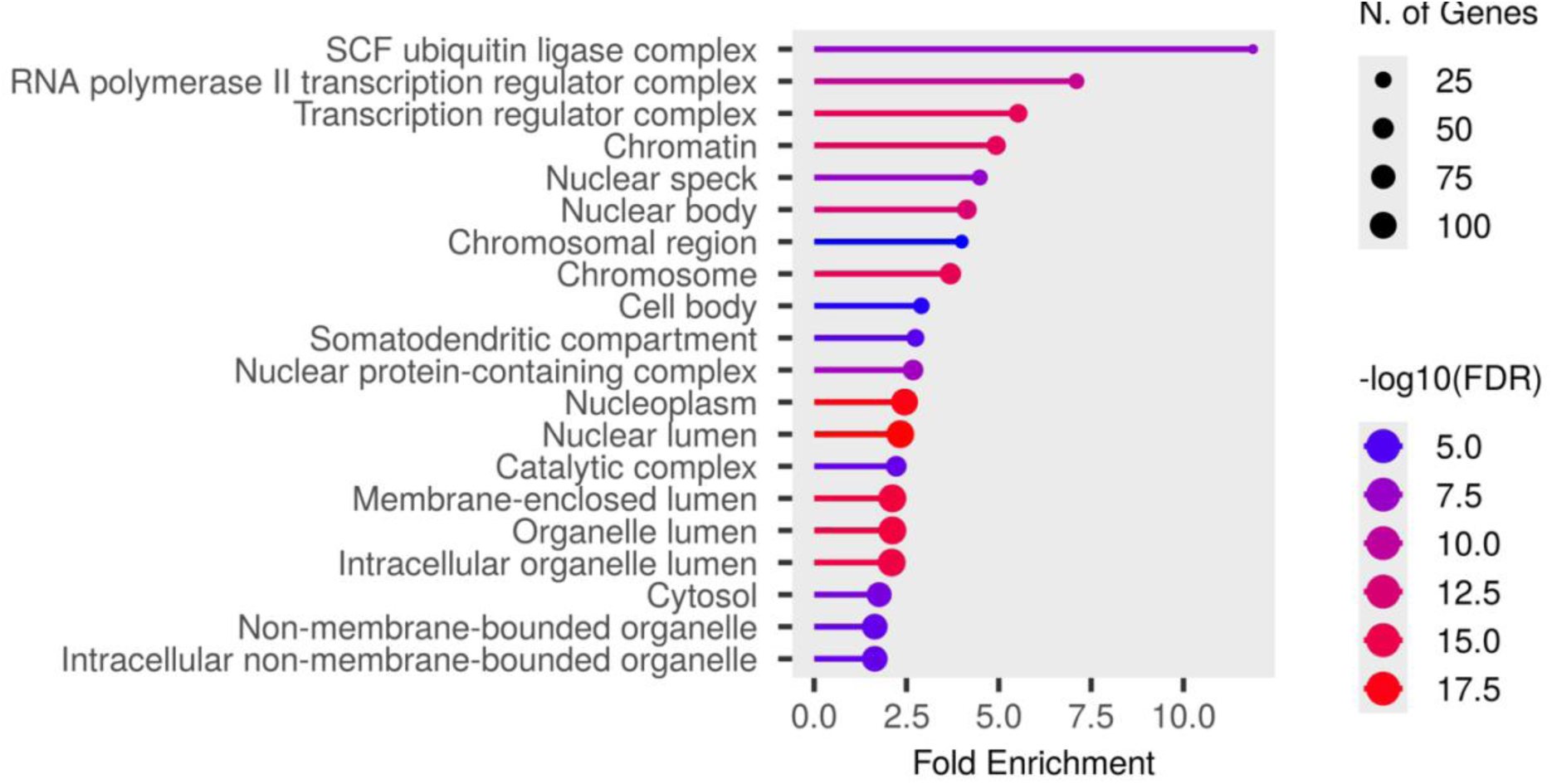
Gene Ontology (GO) Analysis of Cellular Components in NTG-induced rat migraine models vs. saline-injected controls.

**Supplementary 1C.**
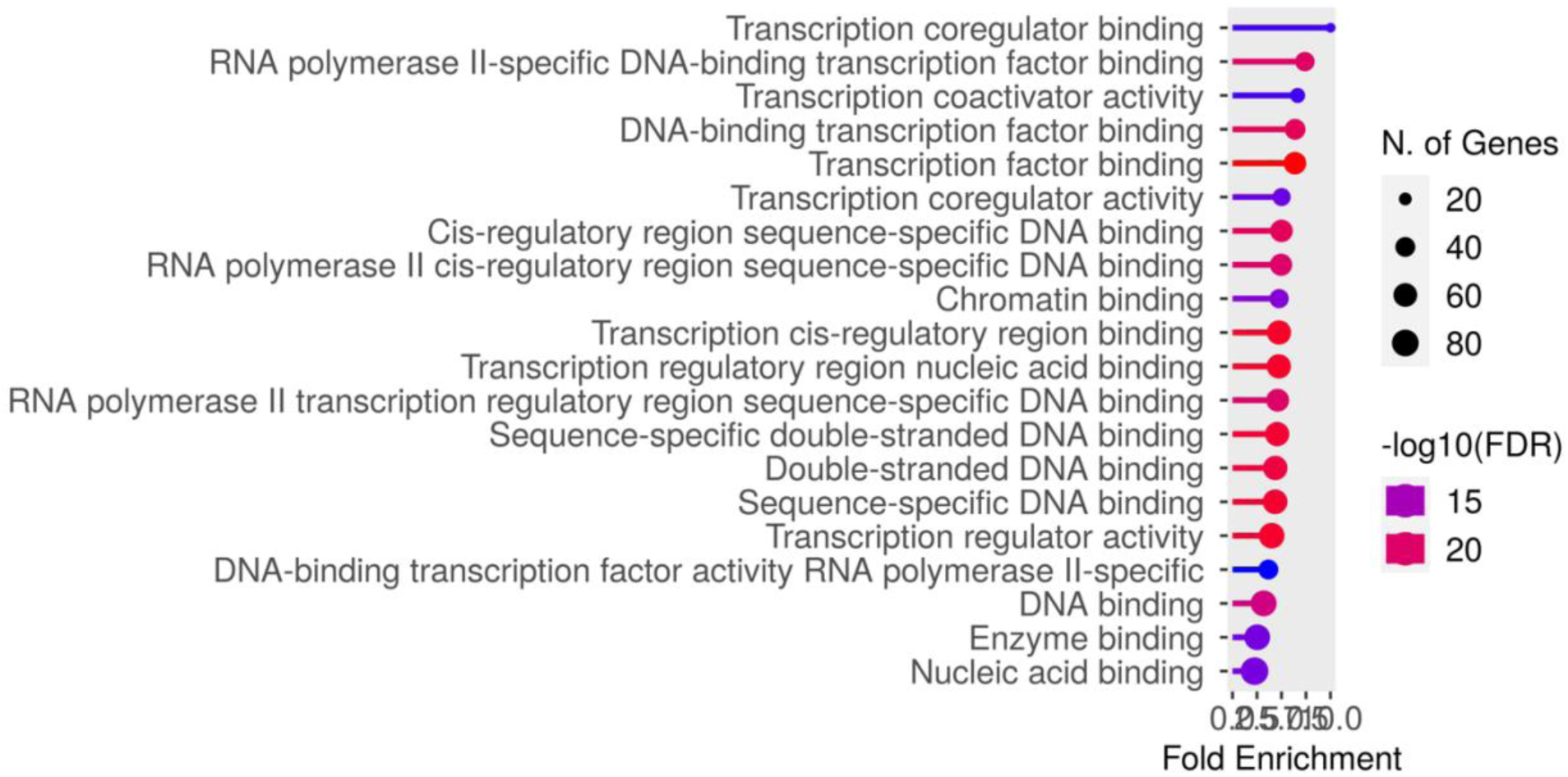
Gene Ontology (GO) Analysis of Molecular Functions in NTG-induced rat migraine models vs. Saline-injected controls.

**Supplementary Figure 1D.**
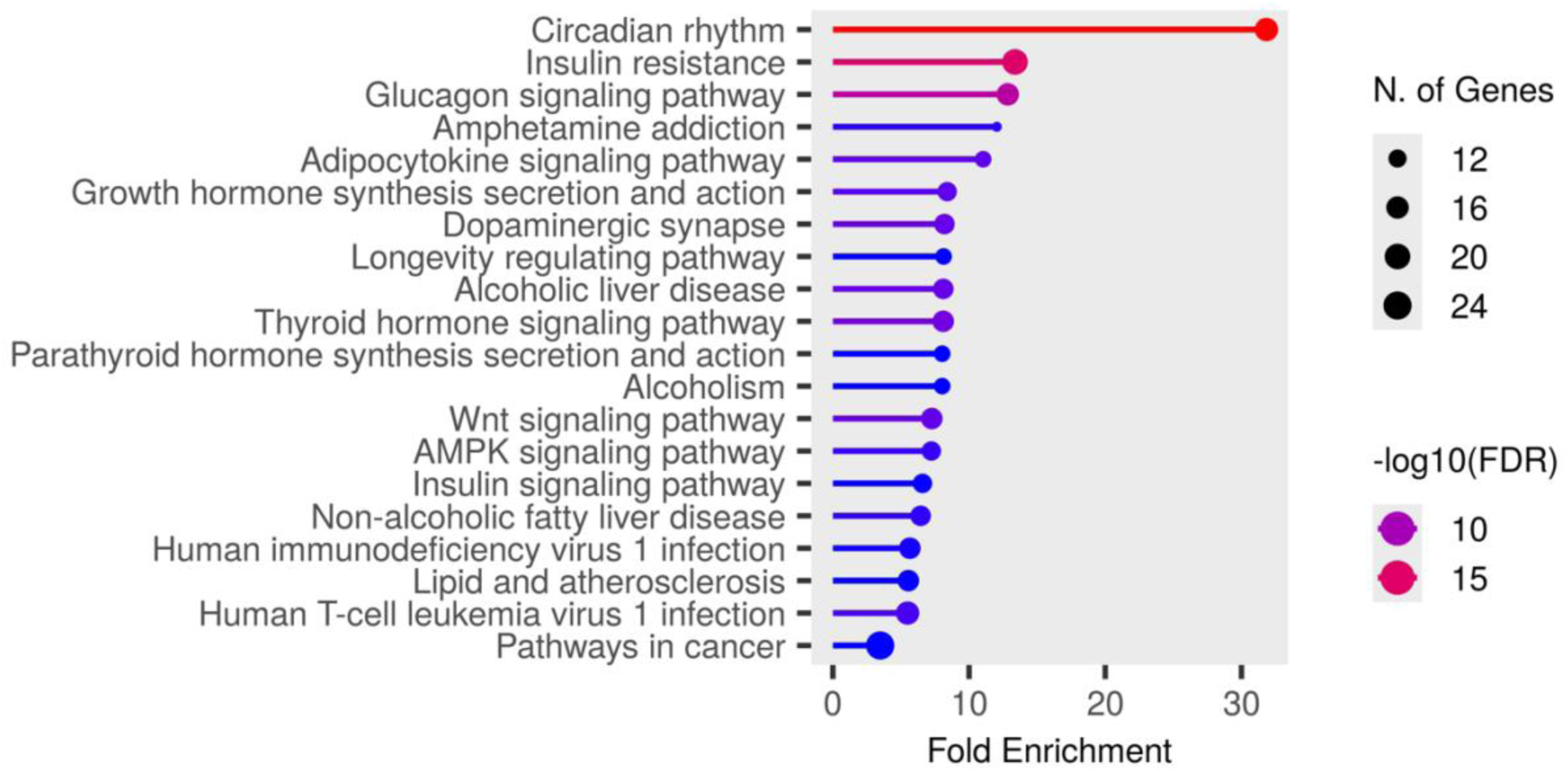
KEGG analysis in NTG-induced rat migraine models vs. Saline-injected controls.

**Supplementary Figure 1E.**
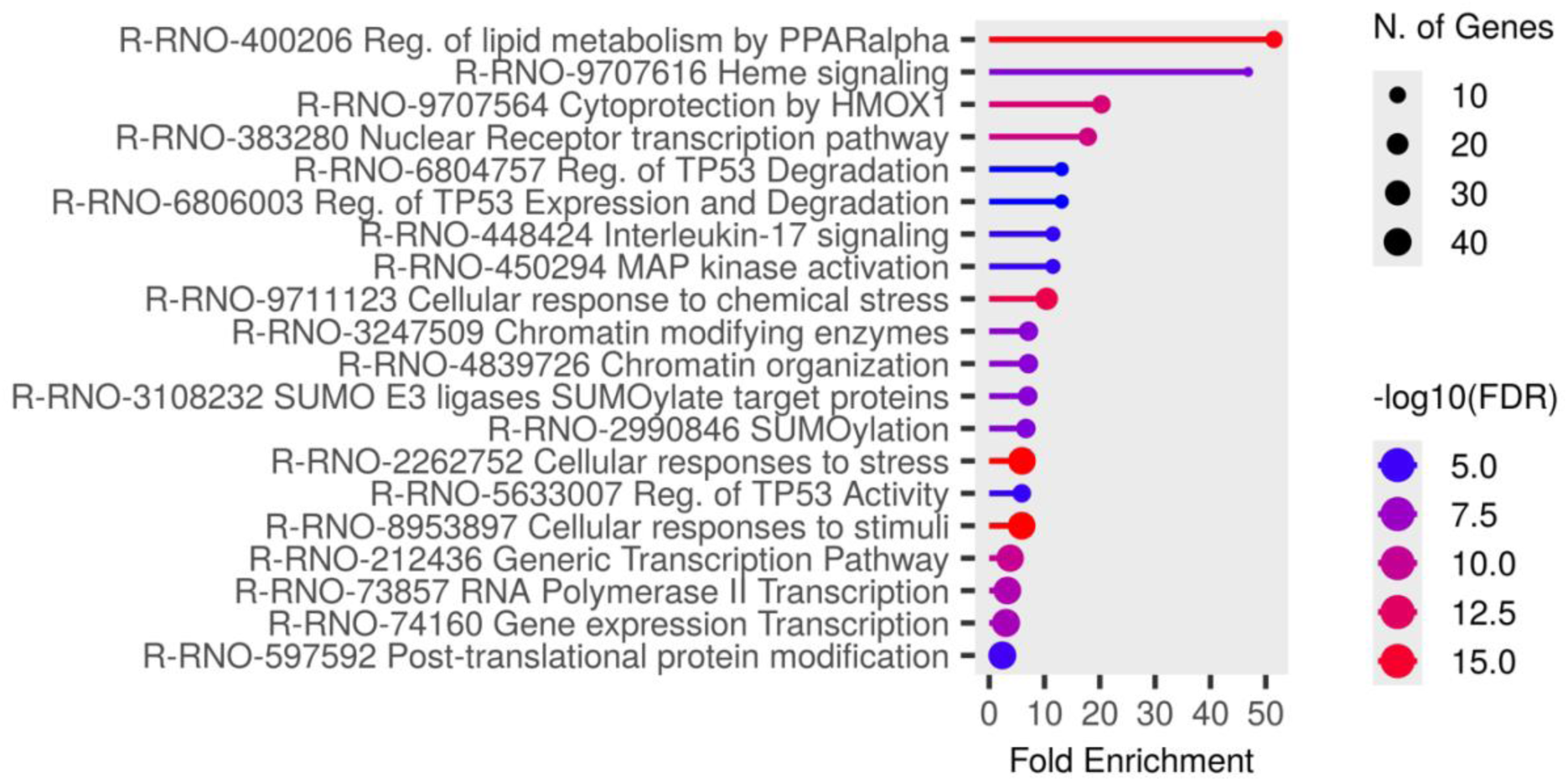
Reactome Enrichment of NTG-induced rat migraine models vs. Saline-injected controls.

**Supplementary Figure 2A.**
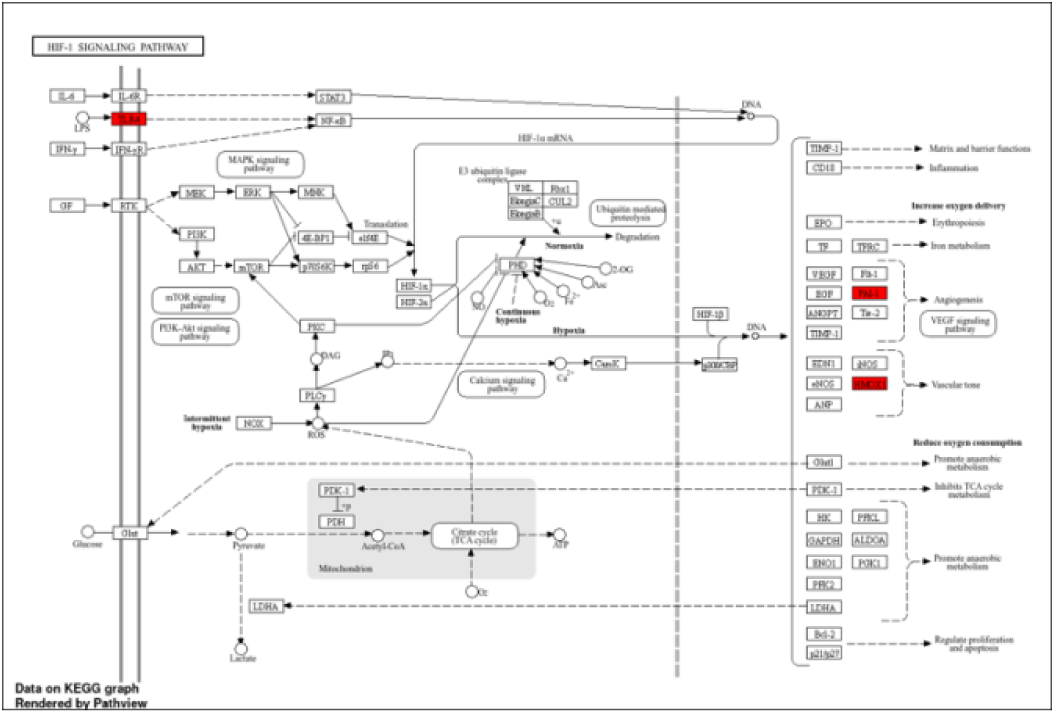
KEGG analysis in Female NTG-induced rat migraine models vs. Female Saline-injected controls.

**Supplementary Figure 2B.**
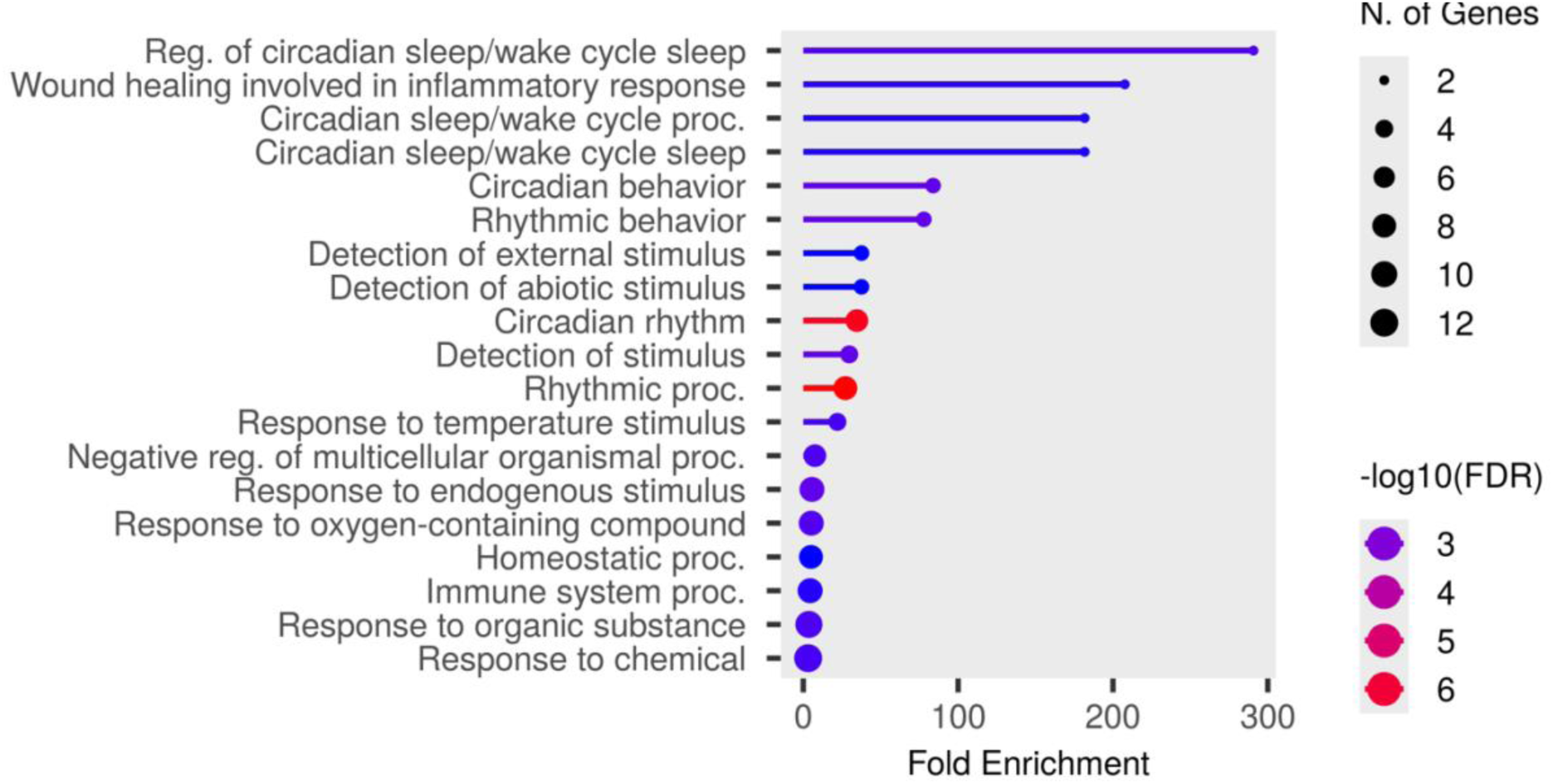
GO Analysis of Biological Processes in NTG-induced Female rat migraine models vs. Female Saline-injected controls.

**Supplementary Figure 2C.**
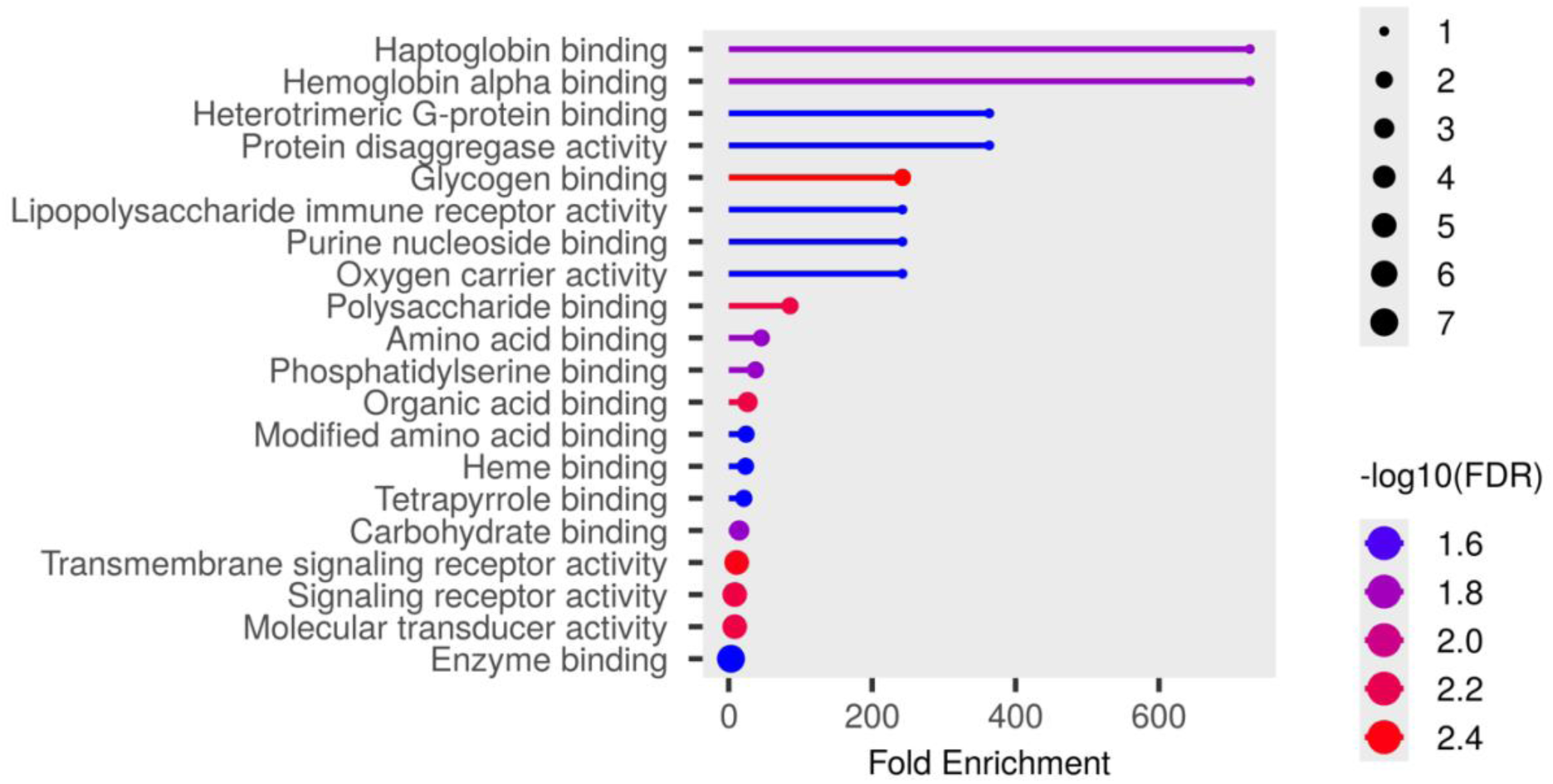
GO Analysis of Molecular Functions in NTG-induced Female rat migraine models vs. Female Saline-injected controls.

**Supplementary Figure 2D.**
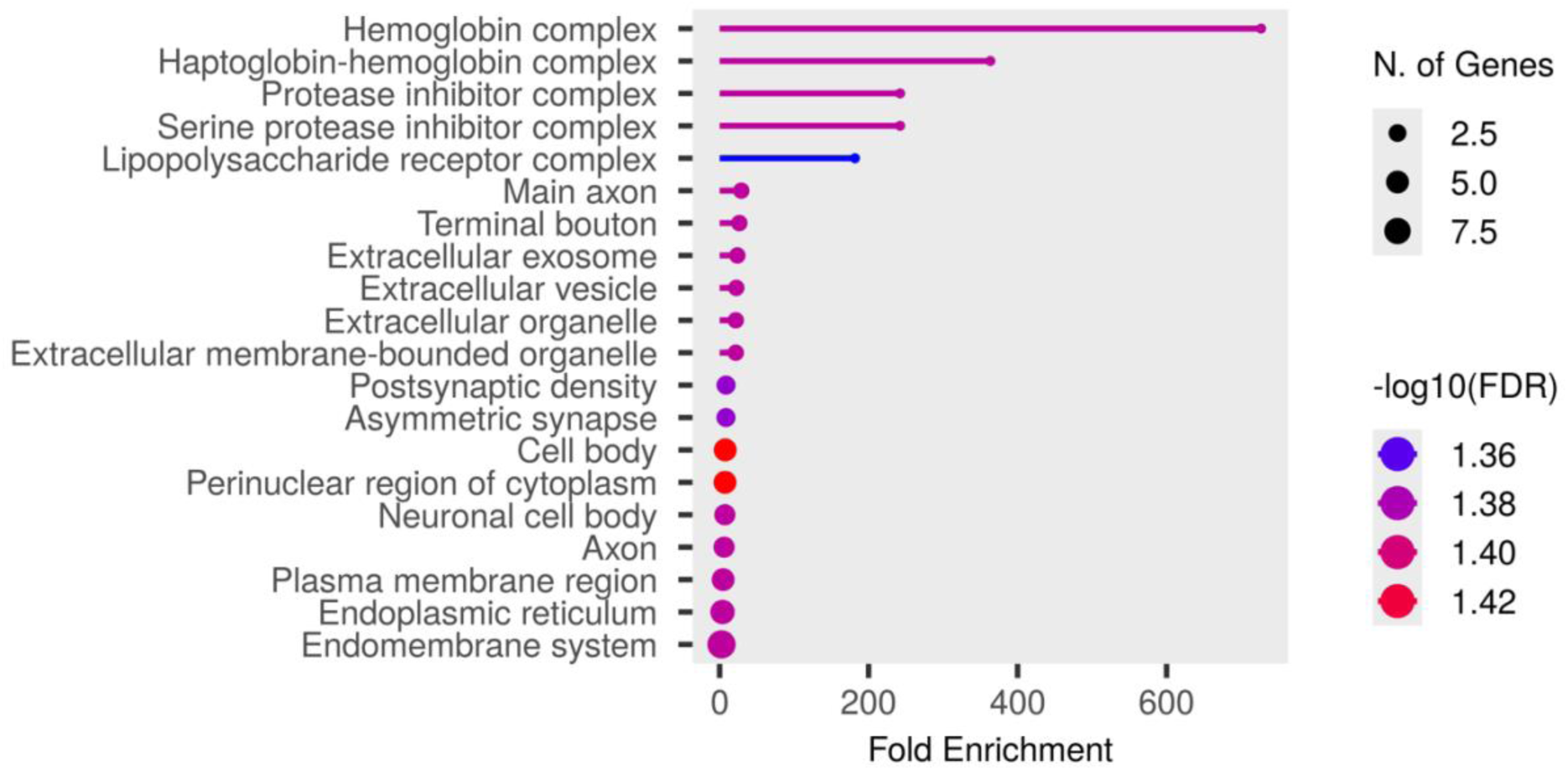
GO Analysis of Cellular Components in NTG-induced Female rat migraine models vs. Female Saline-injected controls.

**Supplementary Figure 3A.**
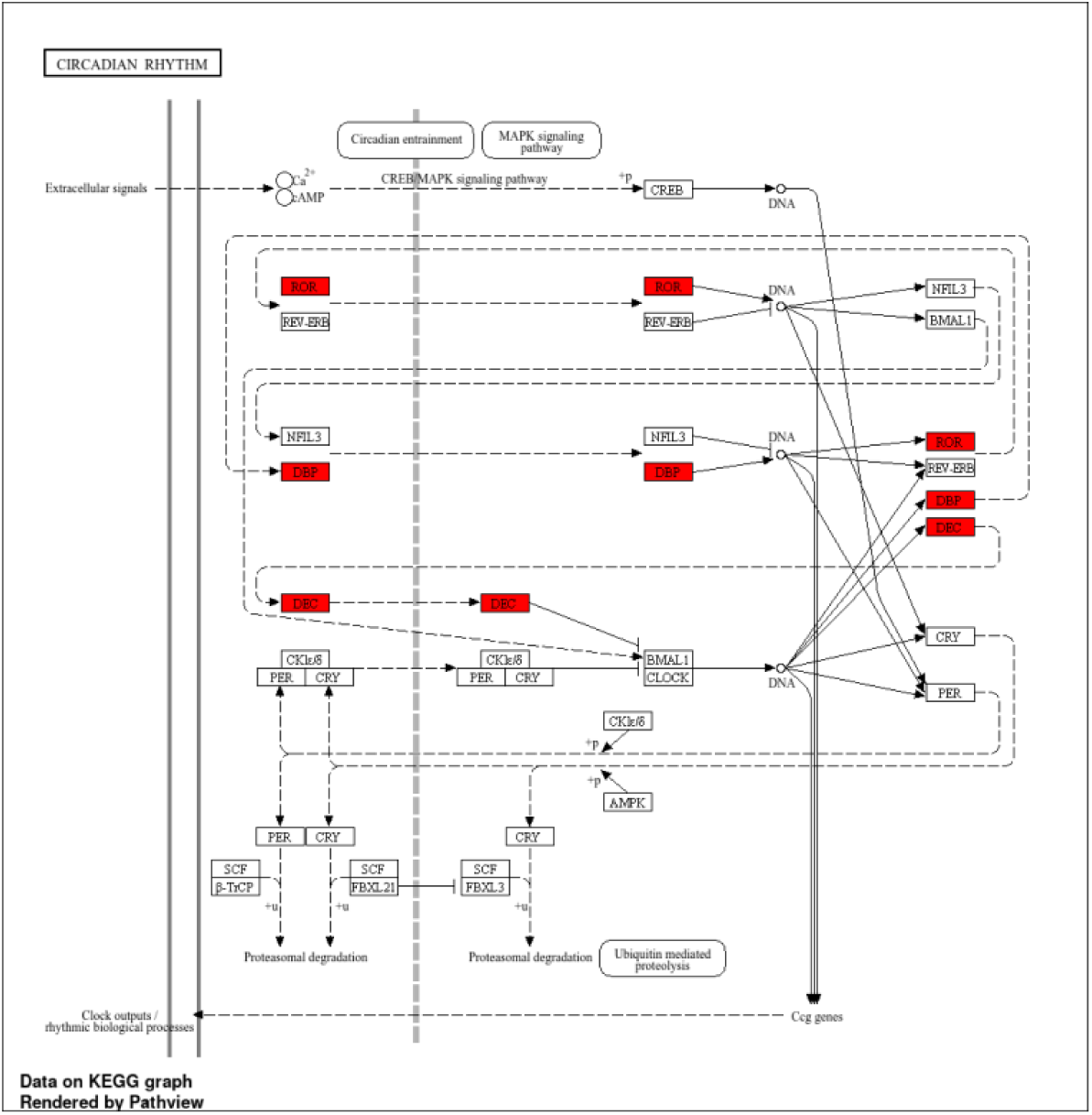
KEGG Analysis in NTG-induced Male rat migraine models vs. Male Saline-injected controls.

**Supplementary Figure 3B.**
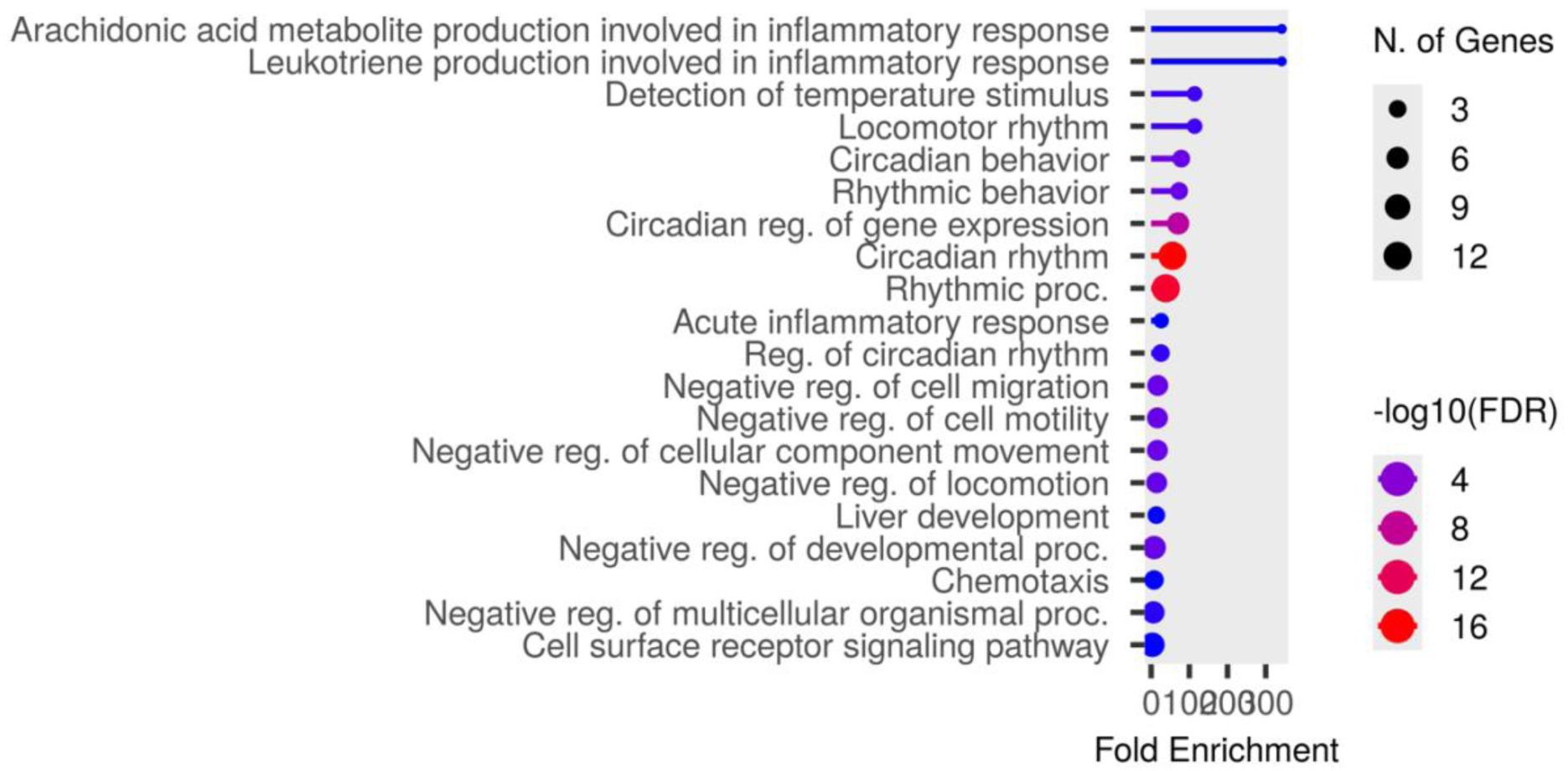
GO Analysis of Biological Processes in NTG-induced Male rat migraine models vs. Male Saline-injected controls.

**Supplementary Figure 3C.**
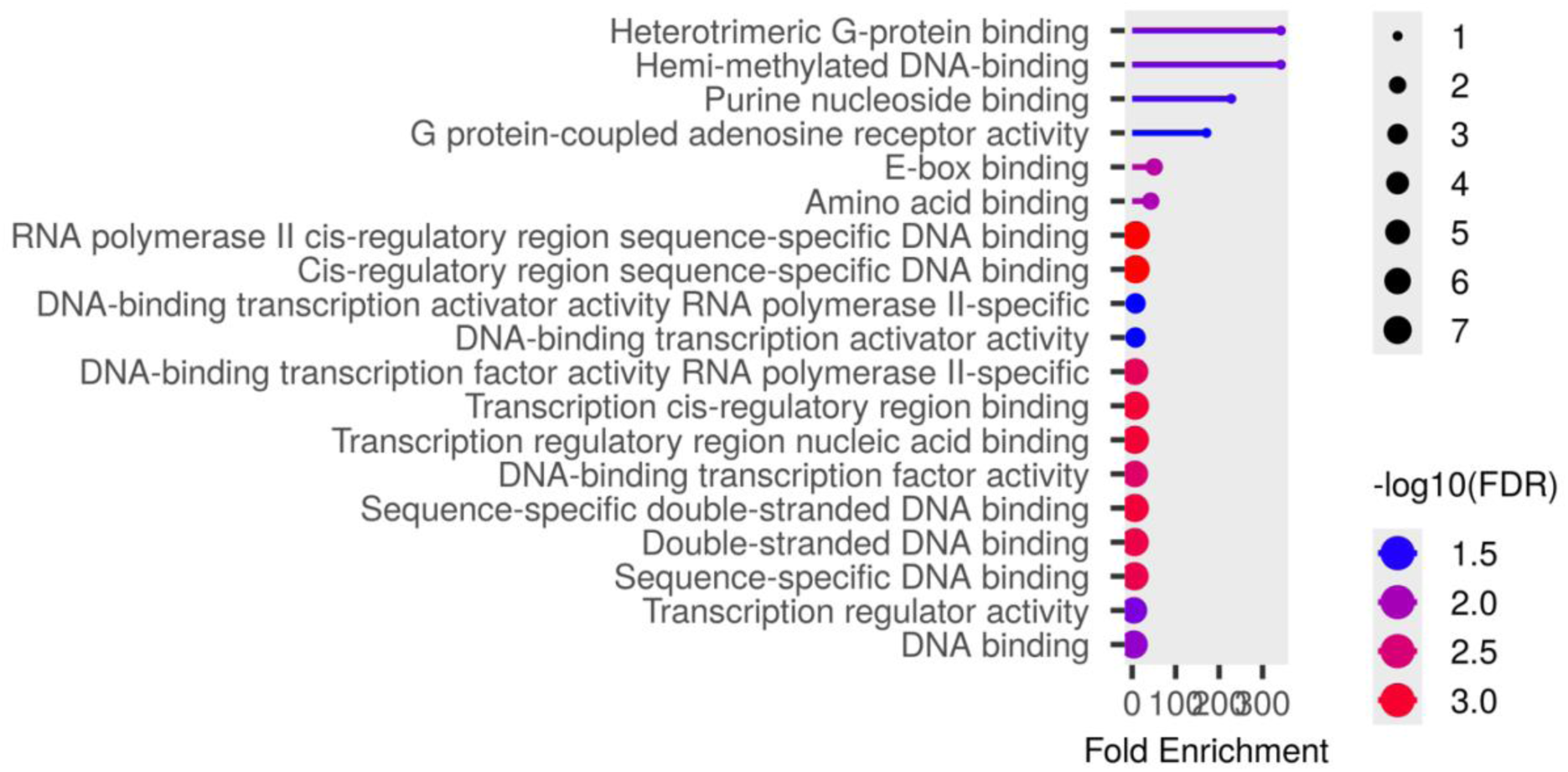
GO Analysis of Molecular Functions in NTG-induced Male rat migraine models vs. Male Saline-injected controls.

